## Supplementary for "Induced neural phase precession through exogeneous electric fields"

*Authors contributed equally

**Supplementary materials**

**Abbreviations**

AC – Alternating current

ACC – Anterior cingulate cortex

AMPA - α-amino-3-hydroxy-5-methyl-4-isoxazolepropionic acid

BA – Brodmann area

DLPFC – Dorsolateral prefrontal cortex

EEG – Electroencephalography

ETP – Educated temporal prediction

FEF – Frontal eye field

GABA_A_ - γ-Aminobutyric acid-A

IN – Inhibitory neuron

M1 – Primary motor cortex

MEP – Motor evoked potential

MPFC – Medial prefrontal cortex

MPM – Medial premotor cortex

OFC – Orbitofrontal cortex

PLV – Phase locking value

PMd – Dorsal premotor cortex

PY – Pyramidal neuron

ROI – Region of interest

tACS – transcranial alternating current stimulation

TMS – Transcranial magnetic stimulation

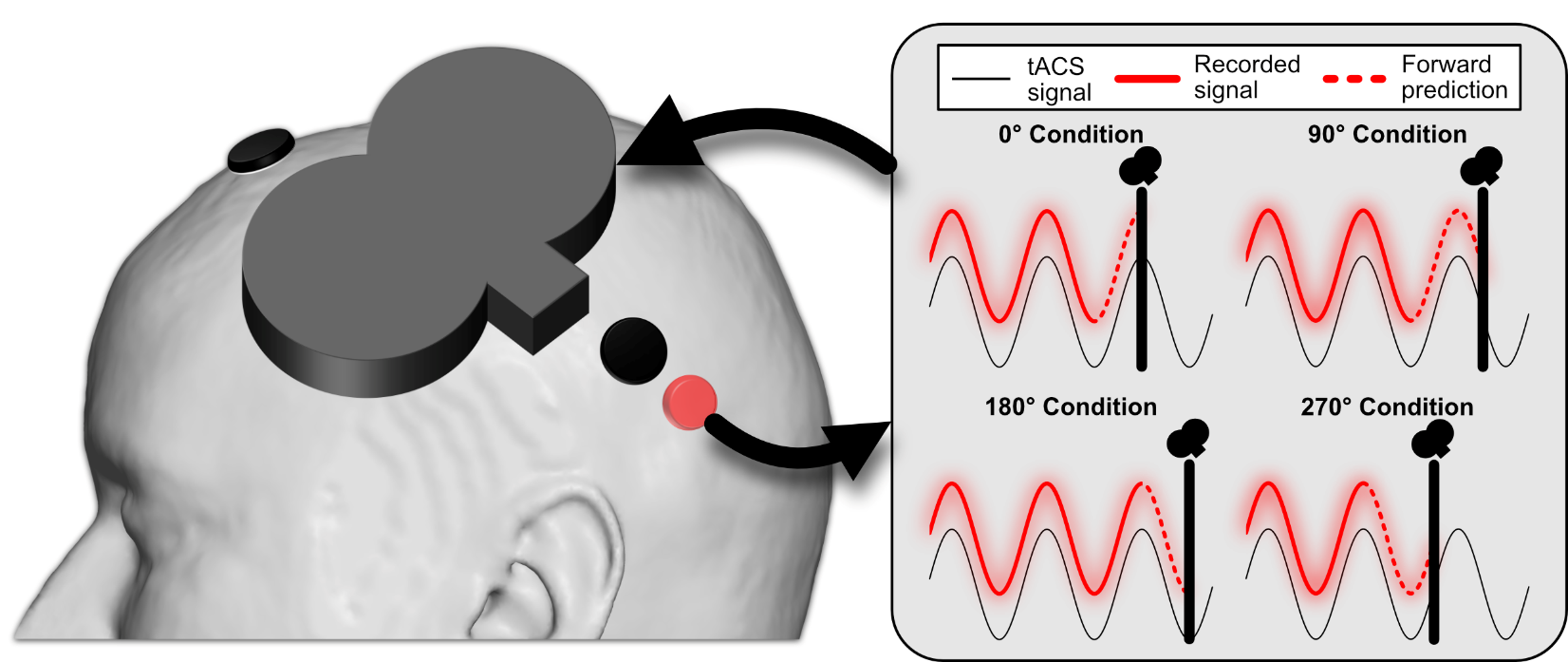

**Supplementary Fig. S1.** Closed-loop tACS-TMS setup. The tACS signal is recorded from the back electrode by a single EEG channel (red). We use our previously validated ETP algorithm for precise phase targeting. Before the experiment the algorithm is trained on 30 seconds of the recorded tACS signal. During real-time tACS-TMS the calculated cycle length is adjusted to inform the forward prediction algorithm to predict upcoming phases (0°, 90°, 180°, 270°). Subsequently a trigger is sent to the TMS apparatus. For details on the ETP algorithm please see^1,2^.

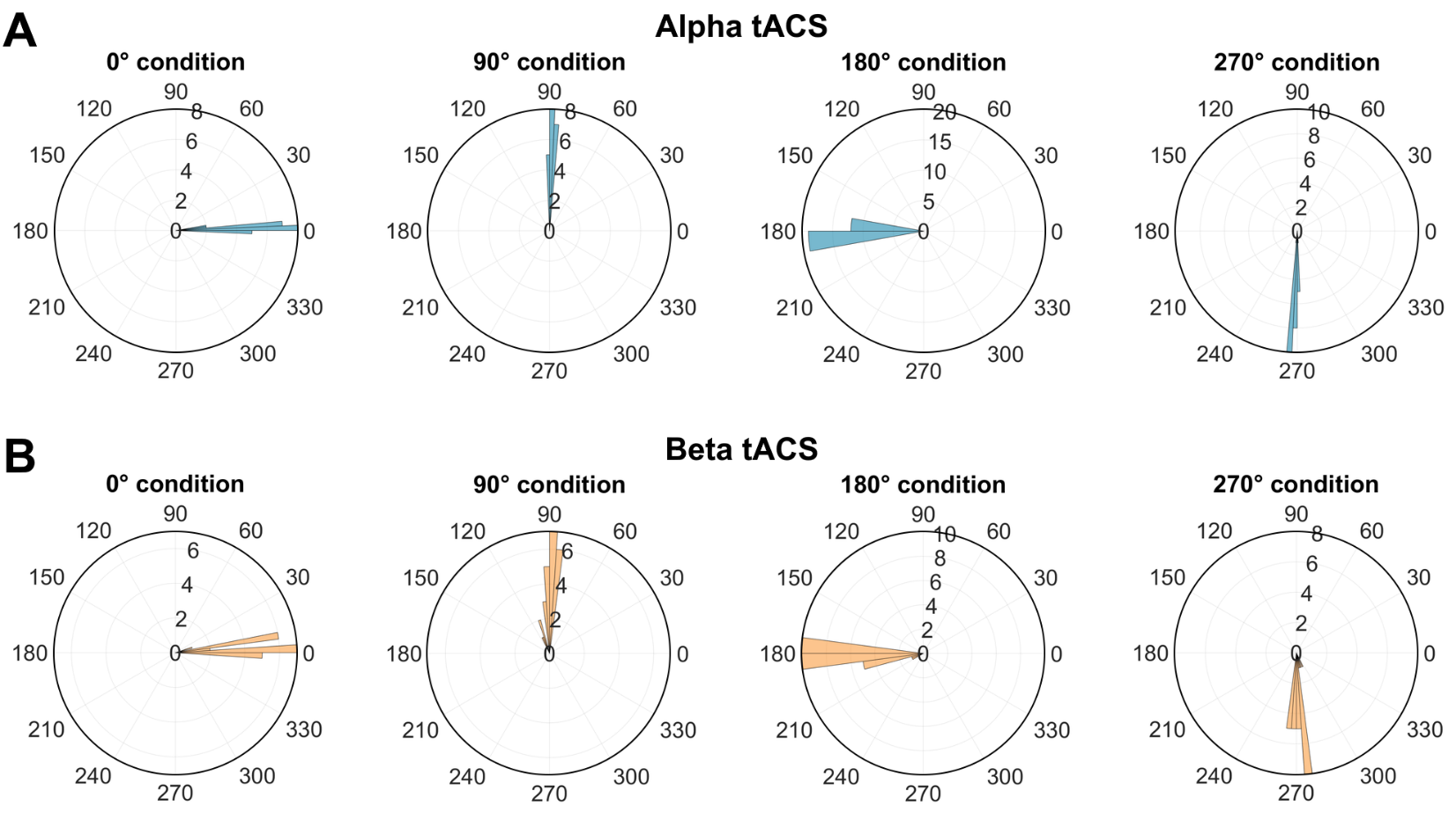

**Supplementary Fig. S2.** Accuracy of phase targeting of our ETP algorithm on the recorded tACS signal for both alpha and beta stimulation. All phases were targeted with high accuracy.

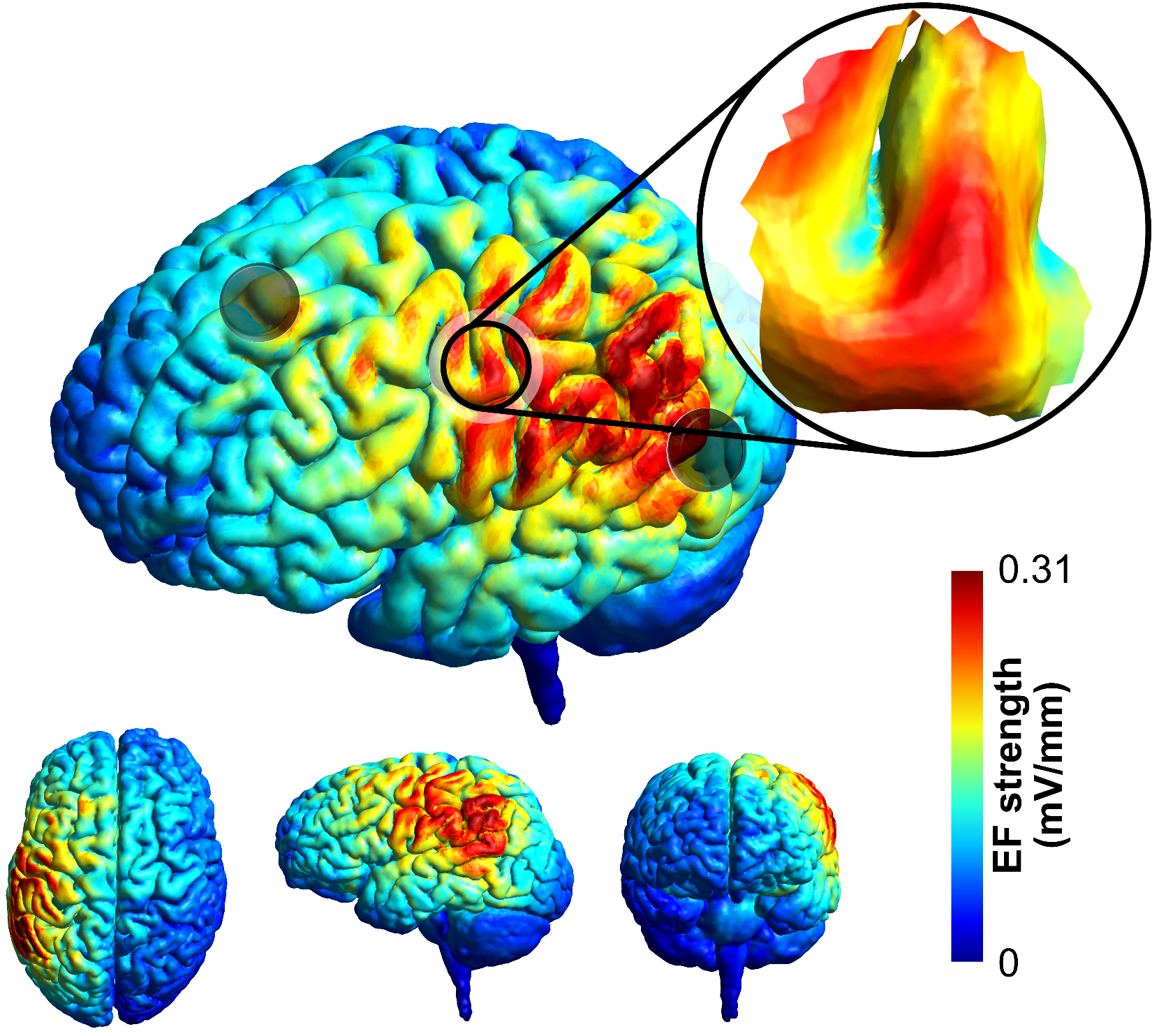

**Supplementary Fig. S3.** Electric field distribution in the human brain using a tACS montage with electrodes placed 7 cm anterior and posterior of the motor hotspot, approximately perpendicular to the orientation of the precentral gyrus.

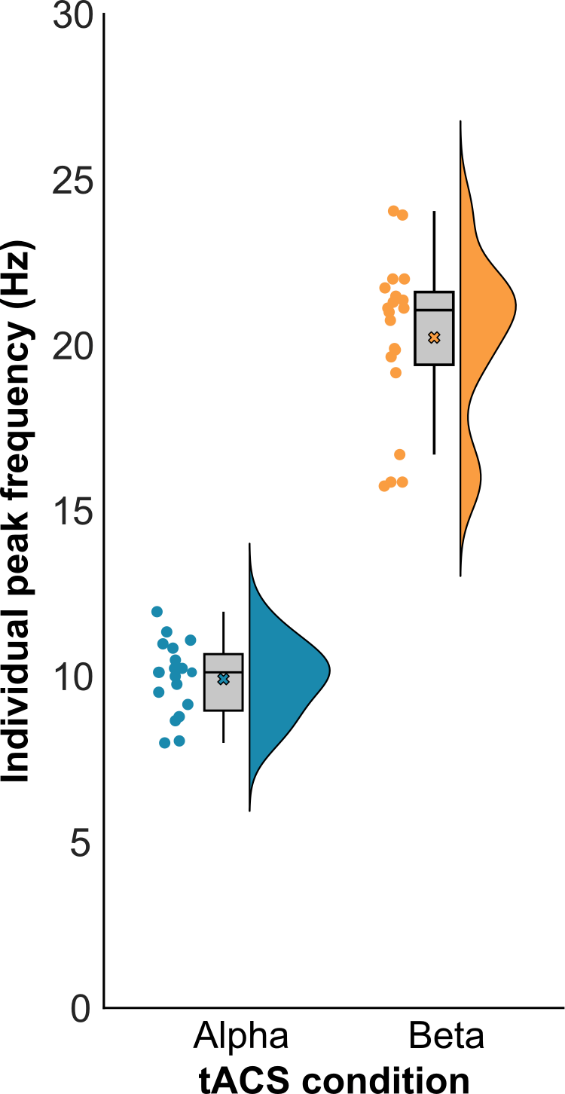

**Supplementary Fig. S4.** Individual peak frequencies of motor cortex alpha and beta frequencies, determined in a 3-minute resting-state EEG at the beginning of each session. These peak frequencies were used for individualized tACS.

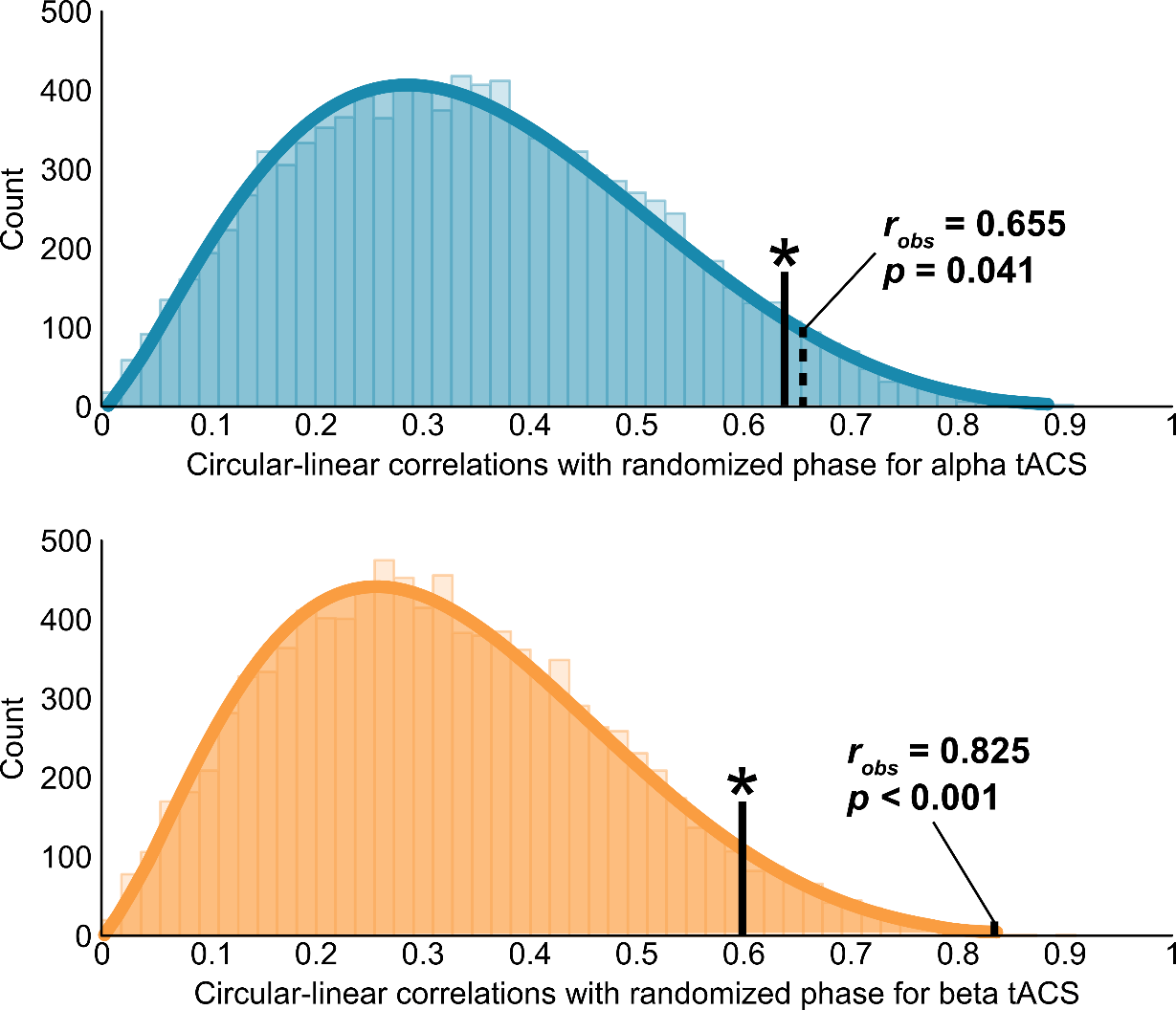

**Supplementary Fig. S5.** Permutation testing on circular-linear correlations in experiment 1. Phase information of MEP was shuffled in 10000 permutations. Actual observed correlation values for alpha (r = 0.655) and beta (r = 0.825) both fall in the 95th percentile of the permutation distribution (p = 0.041 and p < 0.001, respectively), suggesting that the observed phase shifts in human data are unlikely to occur spuriously.

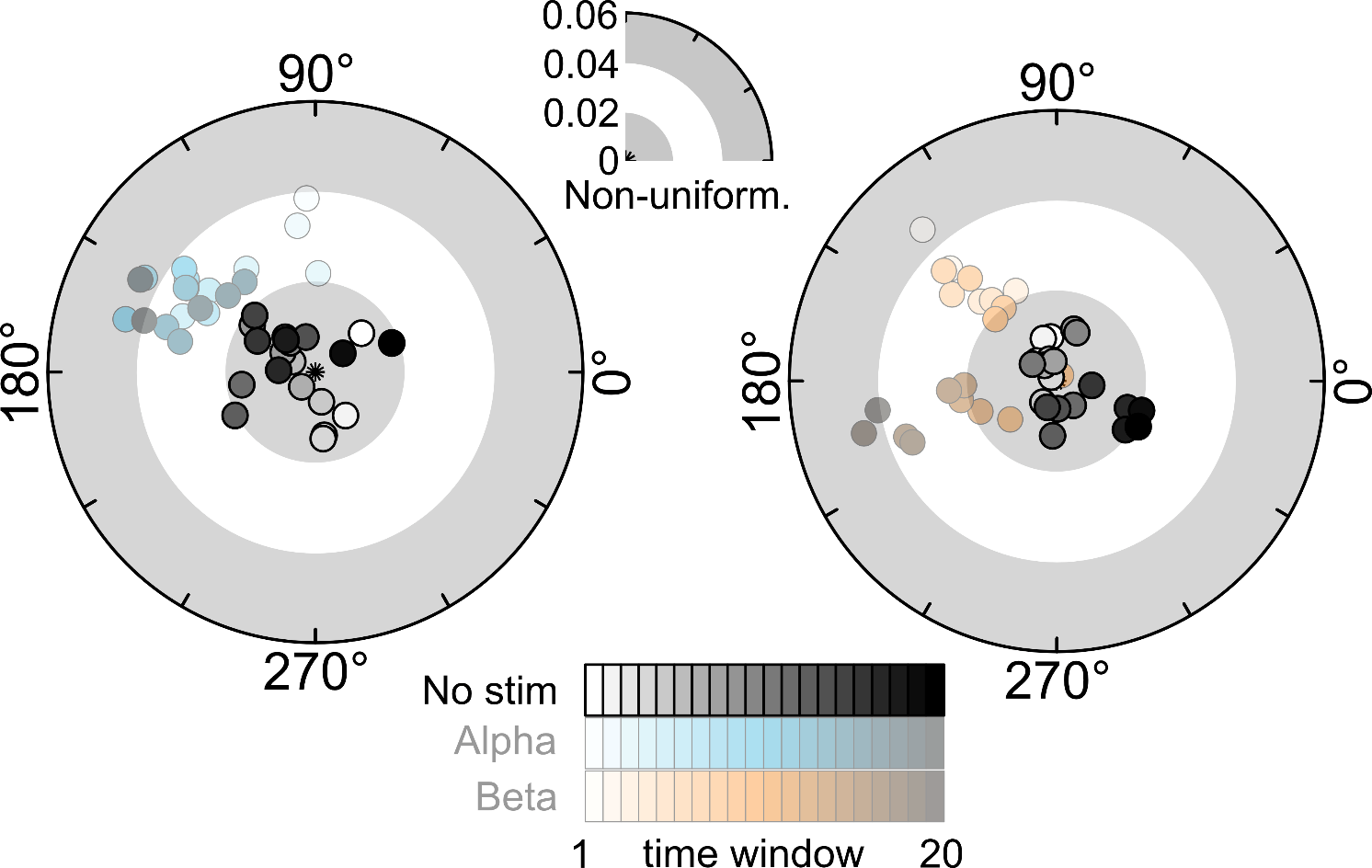

**Supplementary Fig. S6.** Polar plots of changes in phase of primary motor cortex excitability with regards to a virtual tACS signal, without active stimulation. For this control analysis data was taken from Wischnewski et al. (2022)^1^. This dataset follows the same structure as the current experiment, namely 4 blocks of 150 TMS pulses (600 in total), with each block lasting approximately 6 minutes. However, no actual tACS was applied. We applied an alpha and beta virtual tACS signal and extracted the corresponding phases at which the TMS pulse occurred. Subsequently polar vectors were calculated based on phase and MEP amplitude averaged over data per time window, blocks and participants. Each dot represents the polar vector of 55 trials, which equates to approximately two minutes. The sliding window moves in steps of 5 trials (~12 seconds), resulting in 20 windows. On the left the results are shown for the alpha (9.92 Hz) virtual tACS in gray. On the right the results are shown for the beta (20.24 Hz) virtual tACS in gray. Further, the main results with active tACS, as presented in Figure 2 of the main article, are shown transparently for reference. In contrast to active tACS, for the virtual tACS no apparent phase preference, nor phase shift was observed.

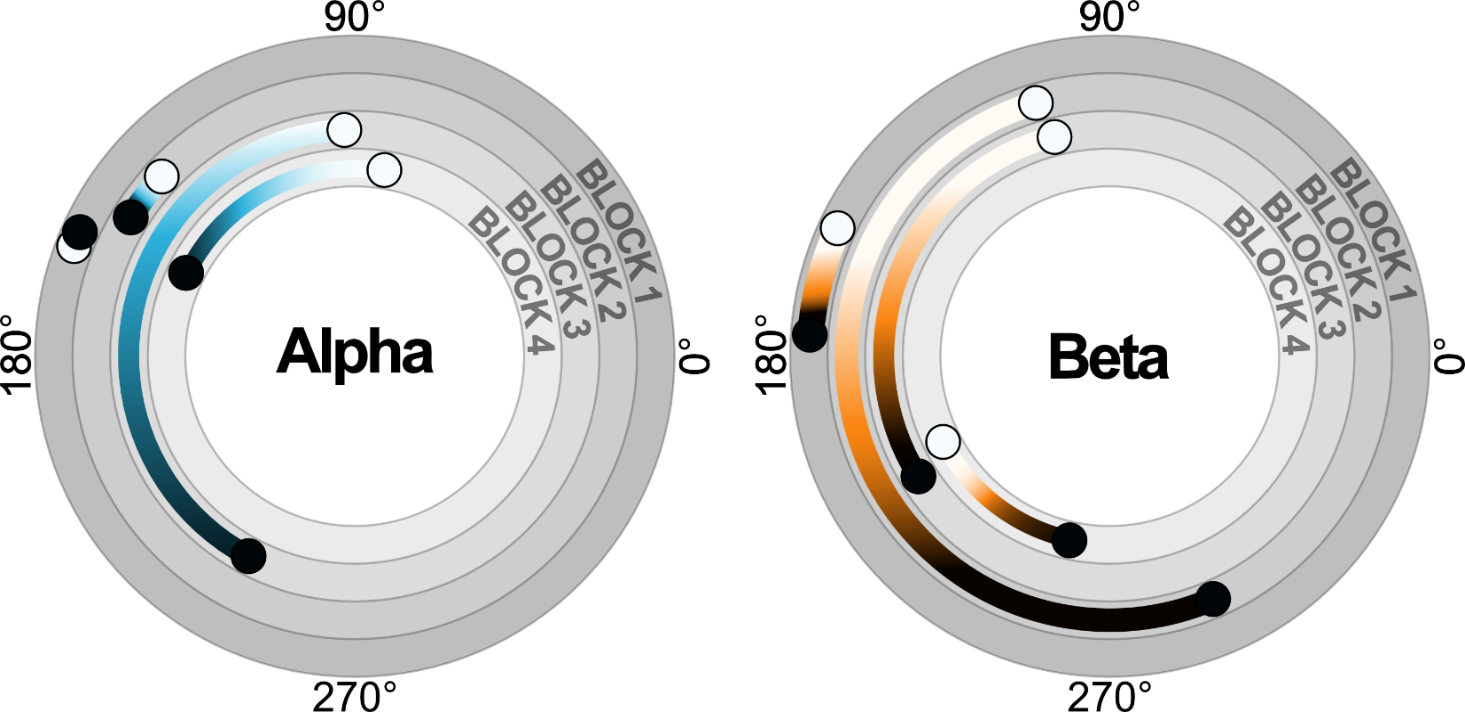

**Supplementary Fig. S7.** Phase shifts of MEPs per stimulation block in experiment 1. Phase of the first and last window are shown. Counter-clockwise phase shifts were observed in 3 out of 4 blocks for alpha tACS, and all blocks for beta tACS. Furthermore, at the beginning of each block the phase resets: For alpha tACS the resets were: block 2-1, 137.3°-155.4° = -18.1°; block 3-2, 93.0°-148.5° = -55.5°; block 4-3, 81.2°-241.6° = -160.4°. For beta tACS the resets were: block 2-1, 106.5°-175.8° = -69.3°; block 3-2, 104.0°-295.1° = -191.1°; block 4-3, 212.0°-207.2° = -4.8°.

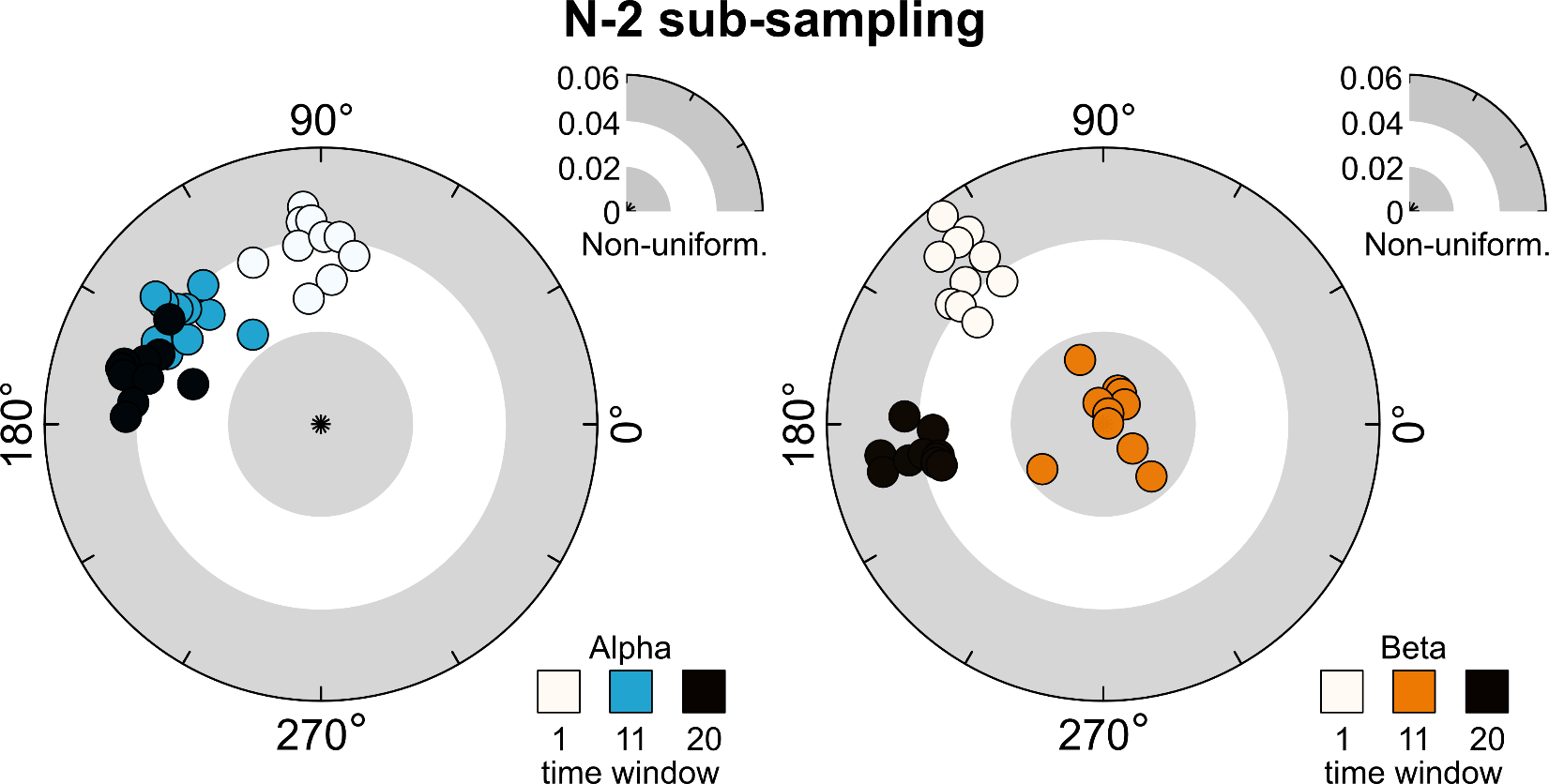

**Supplementary Fig. S8.** Results of an N-2 sub-sampling analysis. This analysis was performed to exclude the possibility that the results of experiment 1 were a consequence of outliers. Specifically, polar plots were generated for 10 sub-samples of n = 18 (leaving out n = 2). Each dot represents one sub-sample. For clarity, we show three of the 20 windows: window 1, 11 and 20. Results were consistent for all sub-samples, for both alpha and beta stimulation, suggesting that the reported main results are not driven by outliers.

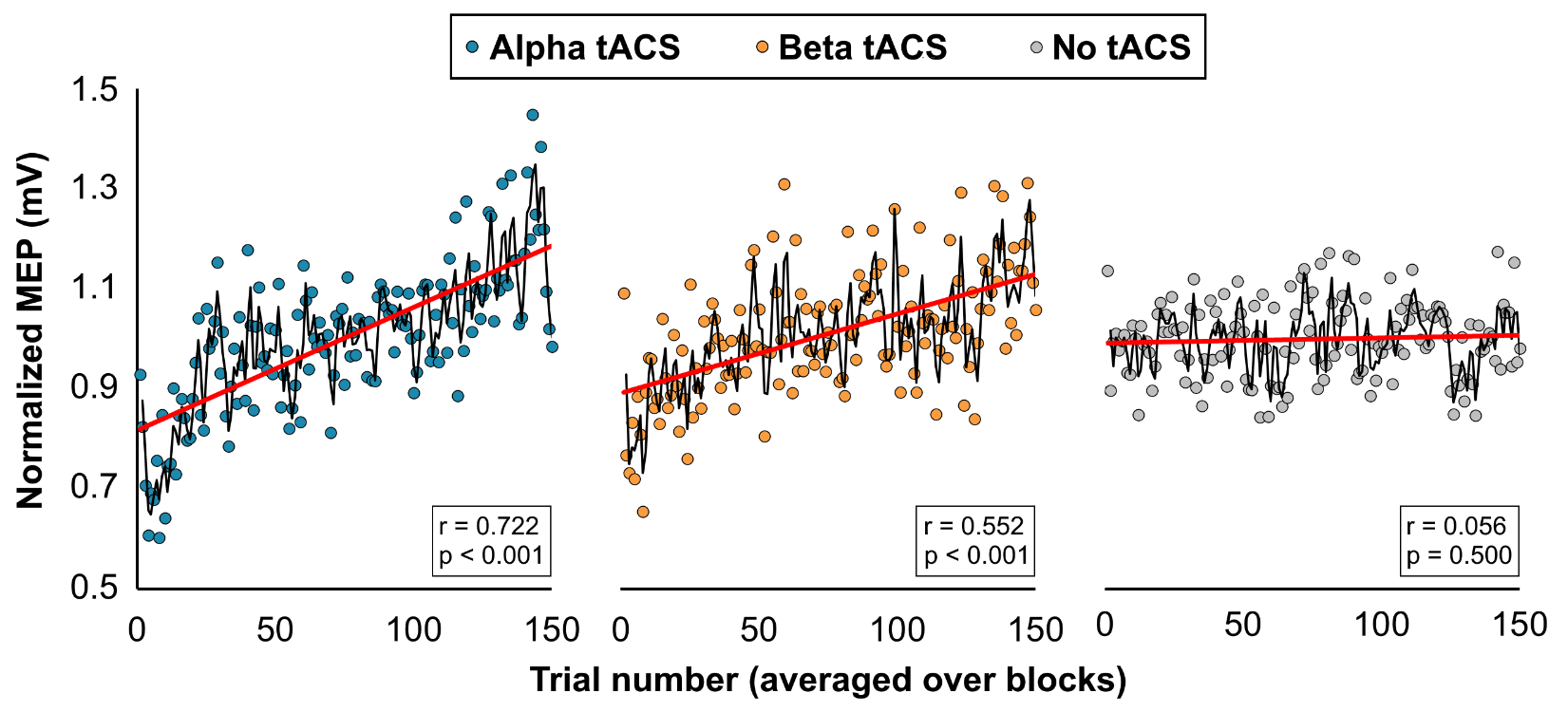

**Supplementary Fig. S9.** Trial-based normalized motor cortical output, expressed as MEPs following a single pulse of TMS. MEPs reflect motor cortico-spinal excitability. Responses were averaged over the four AC stimulation blocks. Black line indicates a moving average, and the red line indicates a linear regression line. During both alpha and beta tACS an increase in MEP size was observed (alpha: r = 0.722, p < 0.001; beta r = 0.552, p < 0.001). However, in a condition without AC stimulation that was otherwise identical, no increase in MEP was observed (r = 0.056, p = 0.500; data taken from^1^).

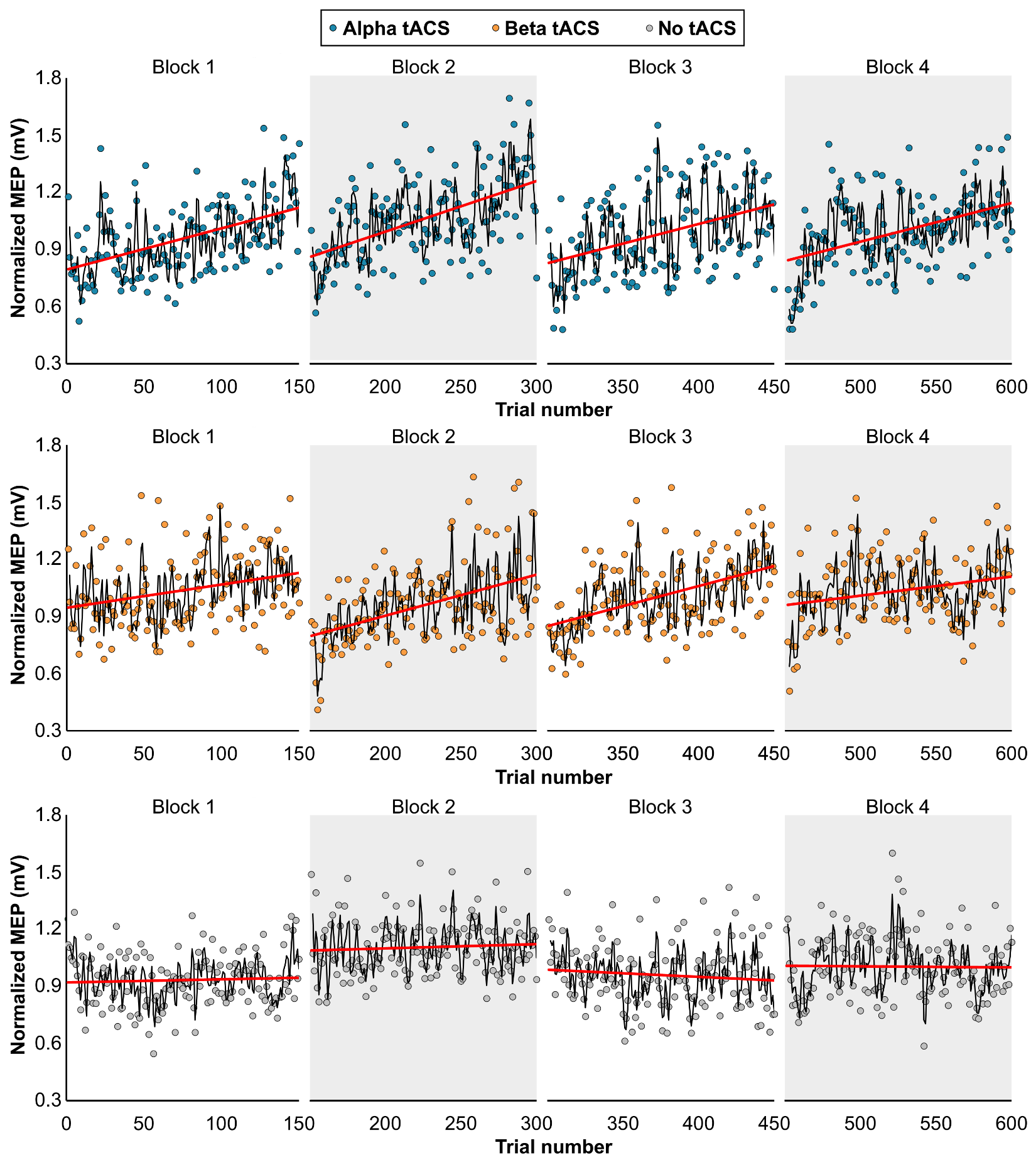

**Supplementary Fig. S10.** Similar data as in Supplementary Fig. 8., but per individual block. During both alpha and beta tACS increases in MEP amplitude were observed in each block. No sustained effect was observed over blocks. In a condition without AC stimulation that was otherwise identical, no increase in MEP was observed (data taken from ^1^).

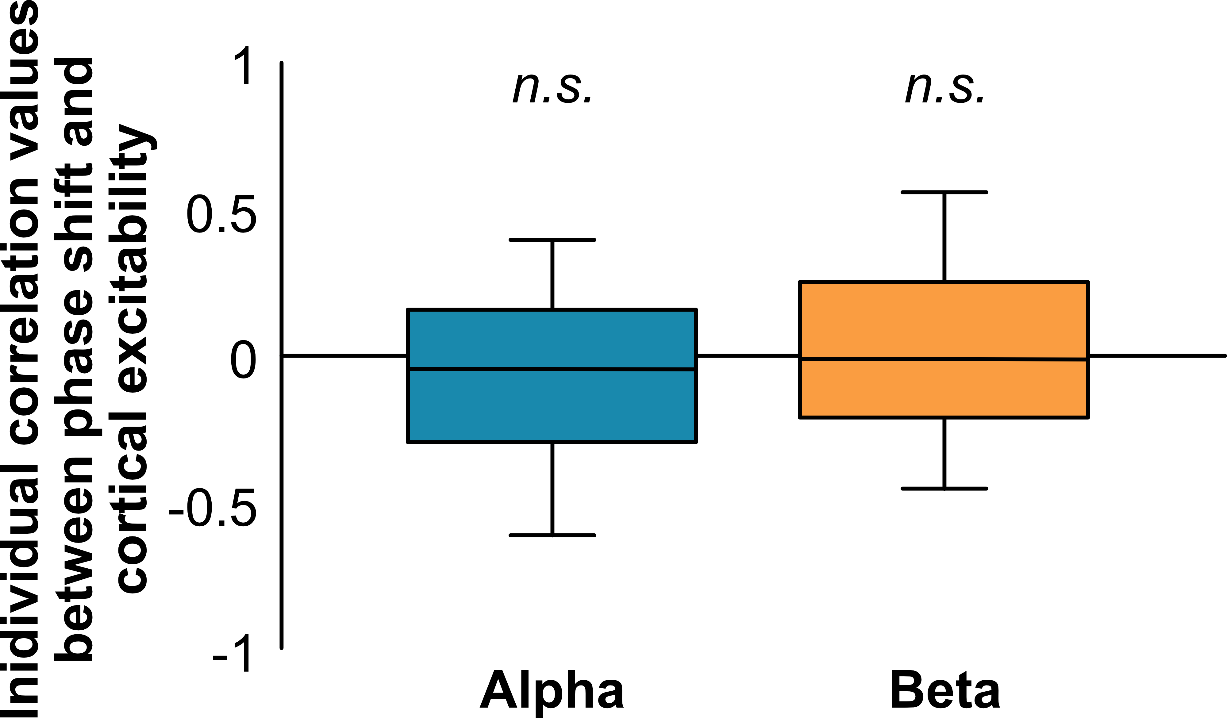

**Supplementary Fig. S11.** Individual correlations (Pearson) between change in cortical excitability and phase shifts. For each participant (n = 20) the window-to-window change of both measures were correlated. On average, r = -0.067 (p = 0.29) for alpha stimulation and r = -0.007 (p = 0.91) for beta stimulation. This suggests that the observed phase shifts (main Fig. 2) and changes in cortical excitability (Supplementary Fig. 8, 9) are independent.

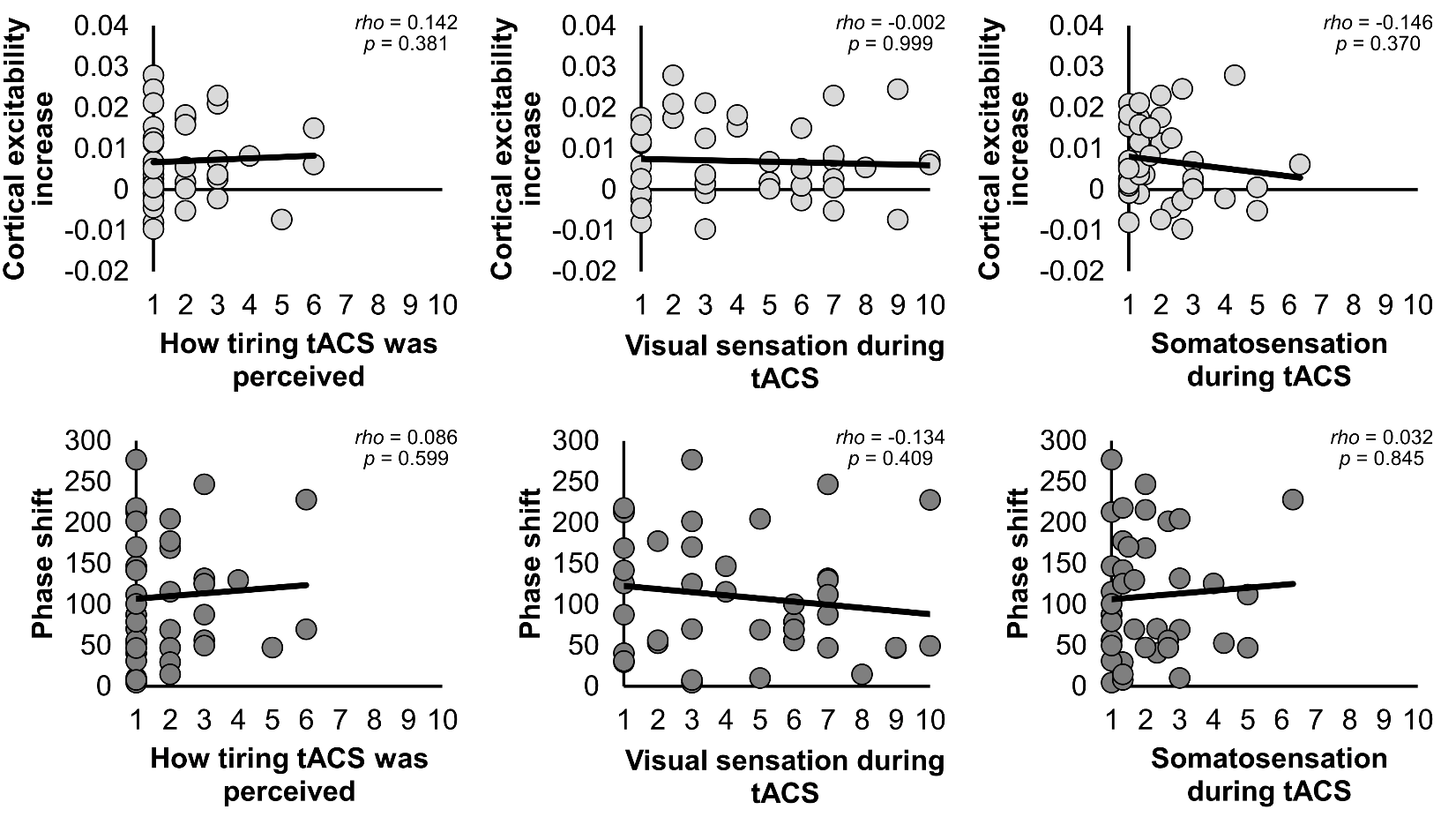

**Supplementary Fig. S12.** Spearman-rank correlations between changes in subjective subject states and cortical excitability changes and phase shifts. Neither visual sensations (phosphenes), nor somatic sensations (skin tingling or itching), nor how tiring the stimulation was perceived was significantly correlated to individual changes in cortical excitability or phase shifts. This suggests that the main results are not confounded by changes in participants’ arousal states.

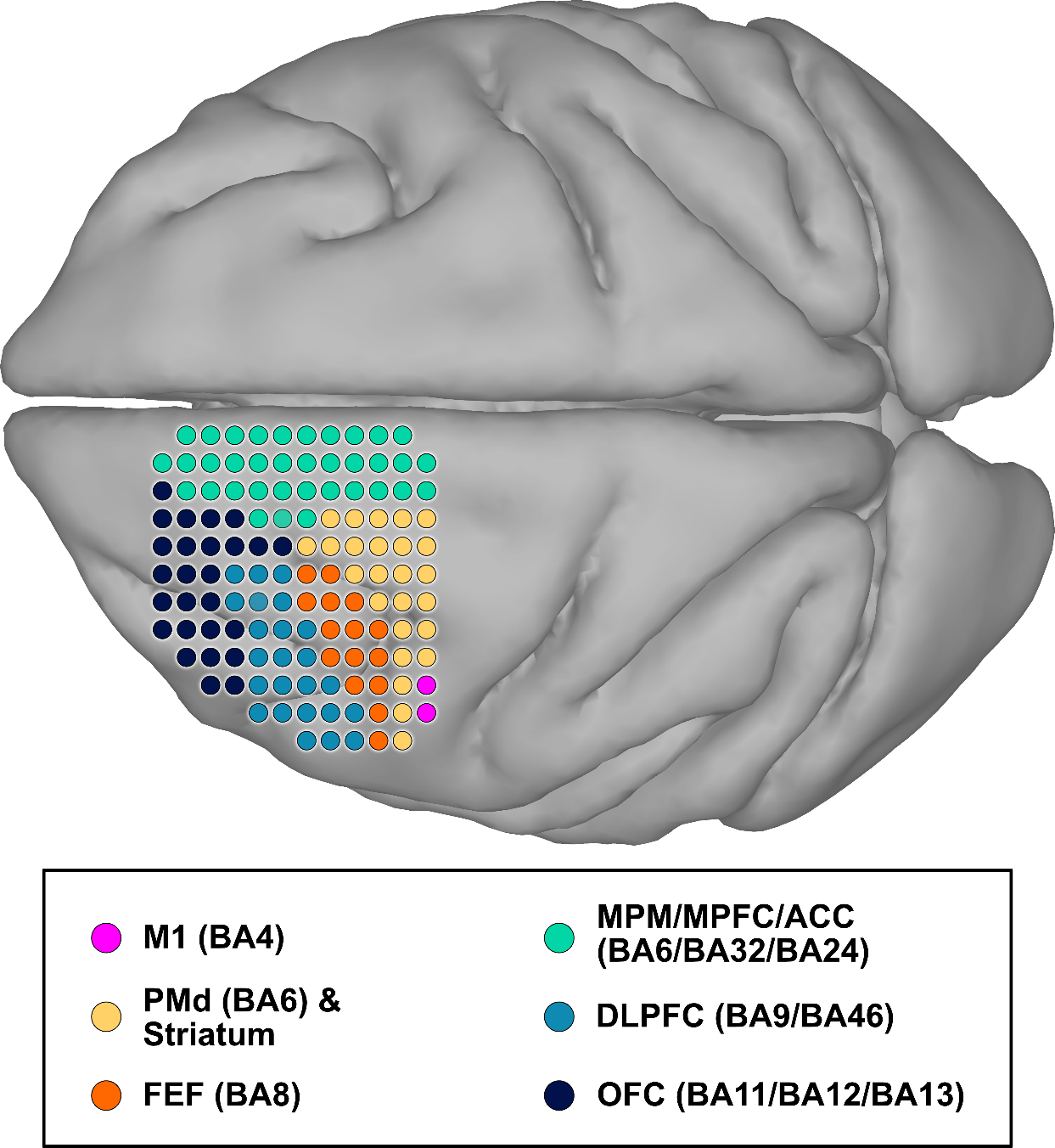

**Supplementary Fig. S13.** Schematic of regional coverage of the 128-channel microdrive recording systems in the NHP. Note that electrode depth differs between electrodes.

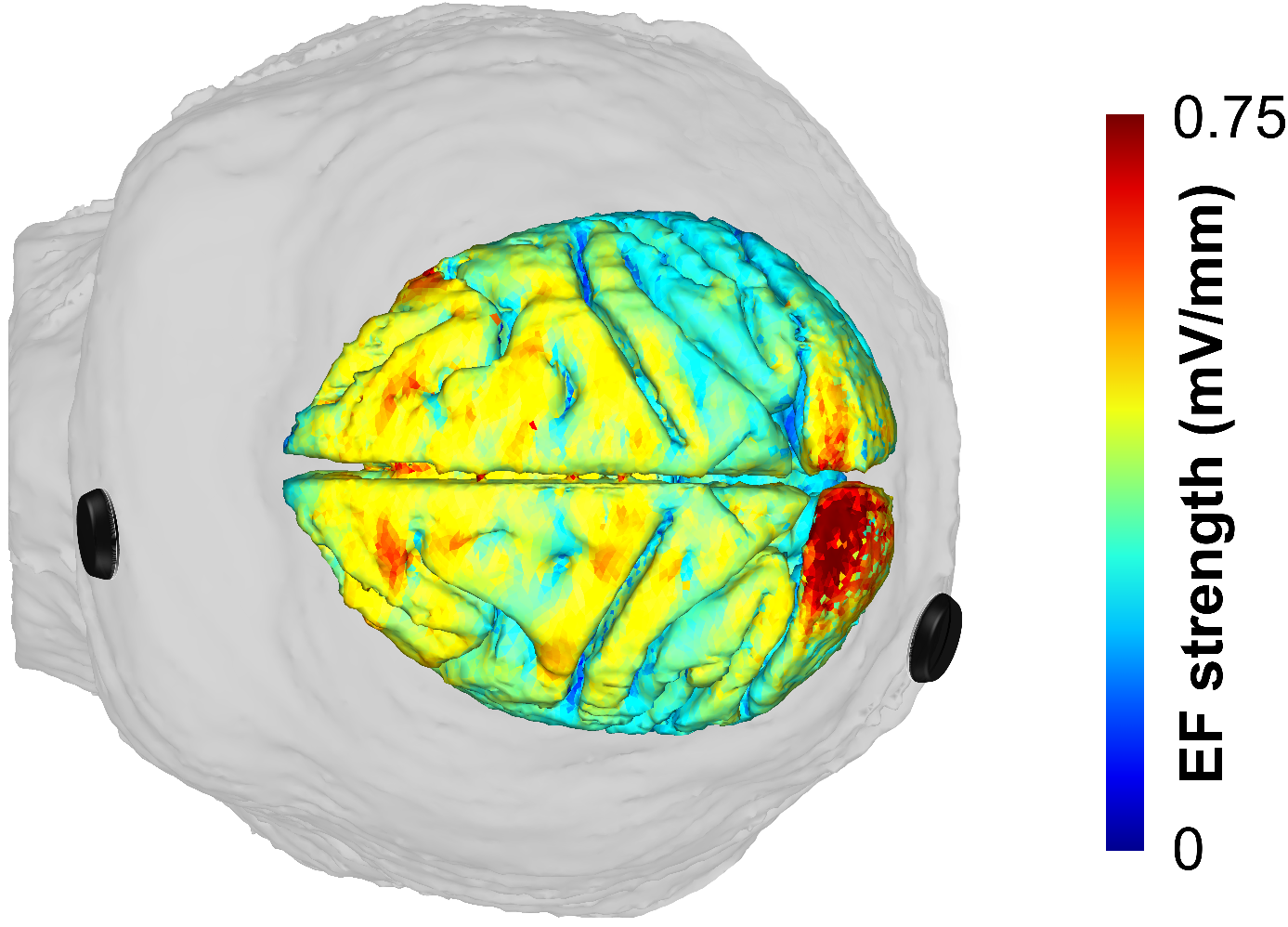

**Supplementary Fig. S14.** Electric field distribution in the NHP brain using a tACS montage with electrodes placed at frontal and parieto-occipital locations (roughly corresponding to FP1 and PO3 in the human 10-20 coordinates).

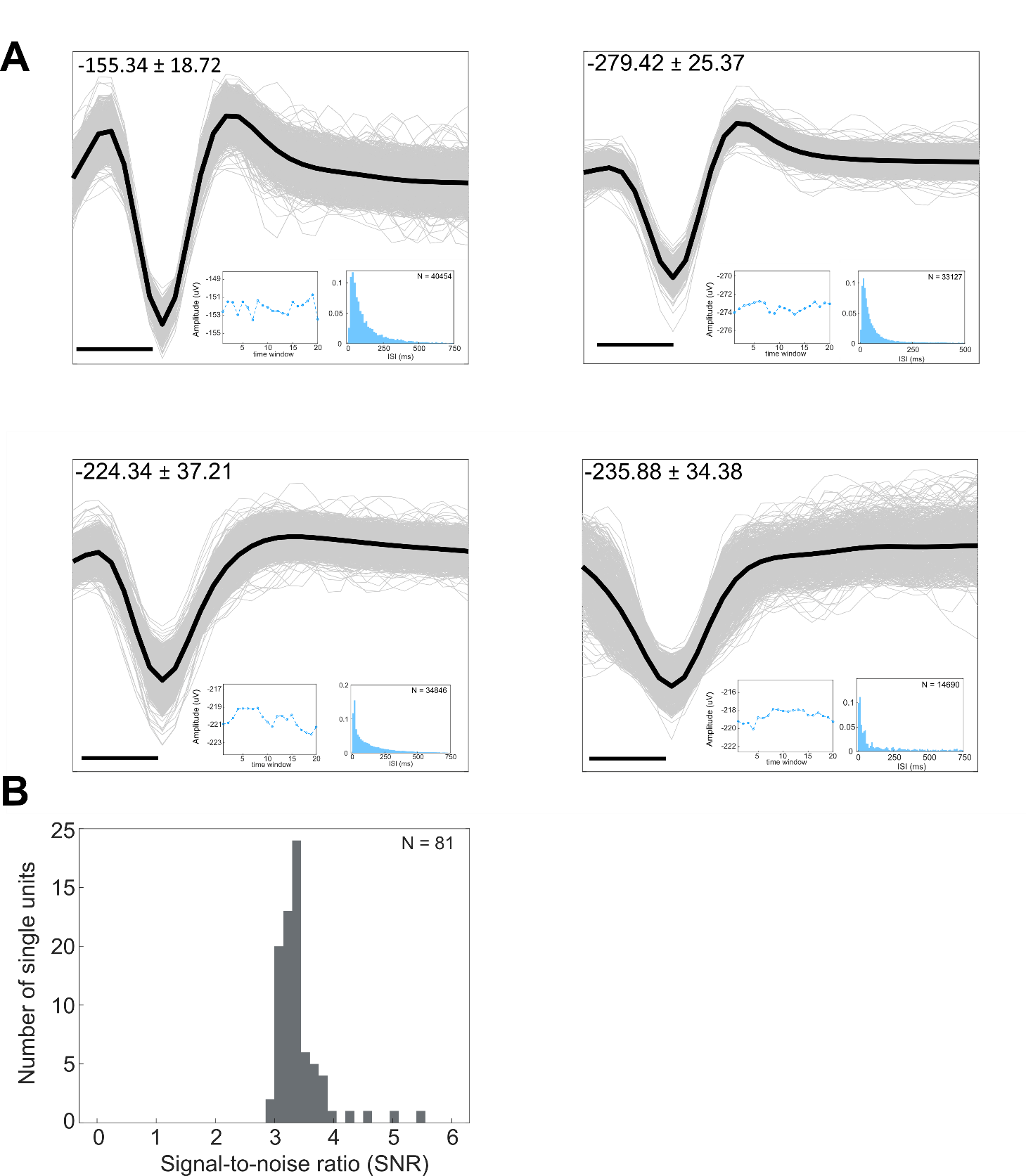

**Supplementary Fig. S15**. Examples of cluster of single units recorded. A) Extracellular spike waveforms of 4 isolated units are shown with additional features. The ISI of the cell is shown in the bottom right with the total number of spikes detected during the recording period. On the left of the ISI is the change in the spike amplitude over the 20 time windows. In the top left corner is the amplitude of the spike (mean ± standard deviation, unit: uV). The thick black line represents the mean average waveform. The black line in the bottom left of each graph represents 200 us. B) Histogram of the signal-to-noise ratio (SNR) of the cells recorded (3.50 ± 0.73, mean ± standard deviation).

**Supplementary Fig. S16**. Schematic of the framework used to quantify neurons behaviors. Once all the neurons were detected, we apply a first threshold to only keep responsive in at least 10 time windows – corresponding to 50% of the total number of time windows (N=20) using a Rayleigh uniformity test. A second threshold was then applied to only keep neurons exhibiting significant circular-linear correlation (p < 0.05). A 3^rd^ final threshold was then applied to keep neurons exhibiting a large phase shift greater than 15° in either direction.

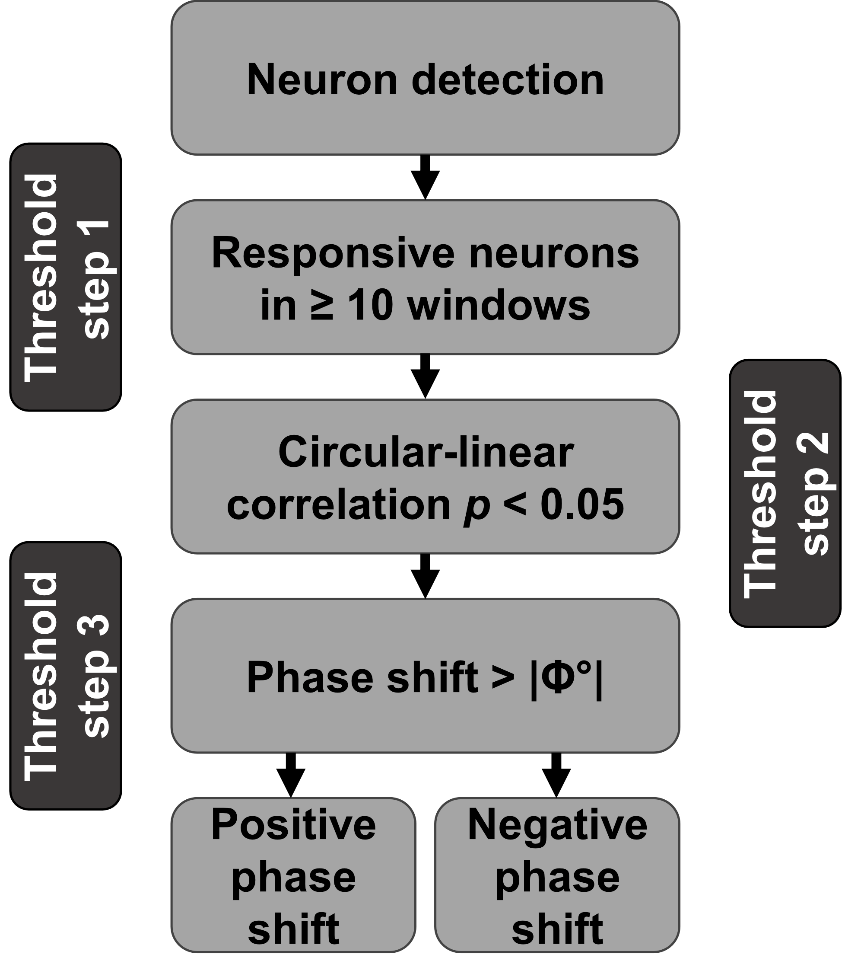

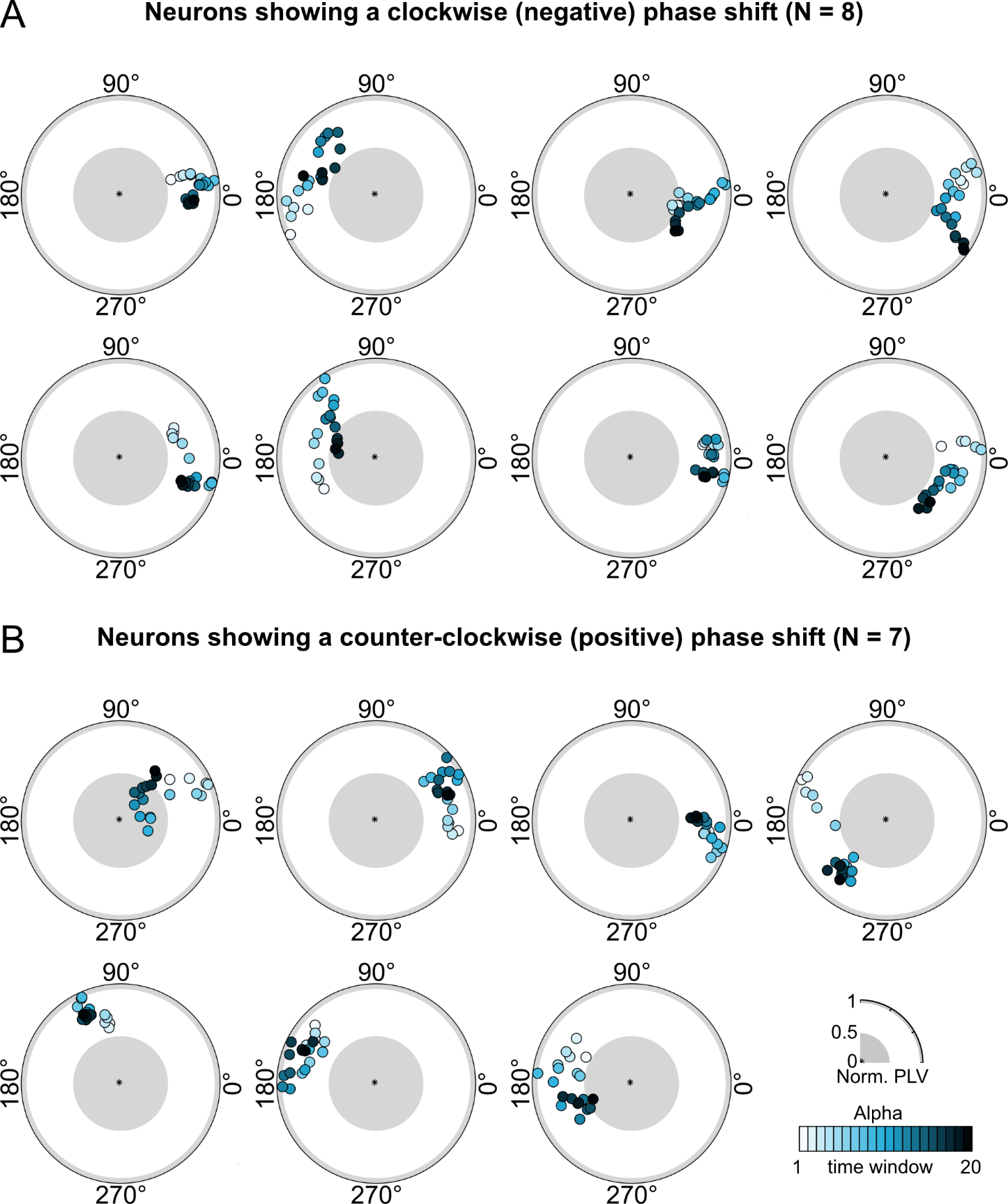

**Supplementary Fig. S17.** Neurons that had a significant circular-linear correlation between time window and preferred phase during alpha AC stimulation. Only neurons with a PLV significantly larger than 0 in at least 10 time windows were selected. A) All neurons (N = 8) that showed phase precession in a clockwise direction. B) All neurons (N = 7) that showed phase precession in a counter-clockwise direction.

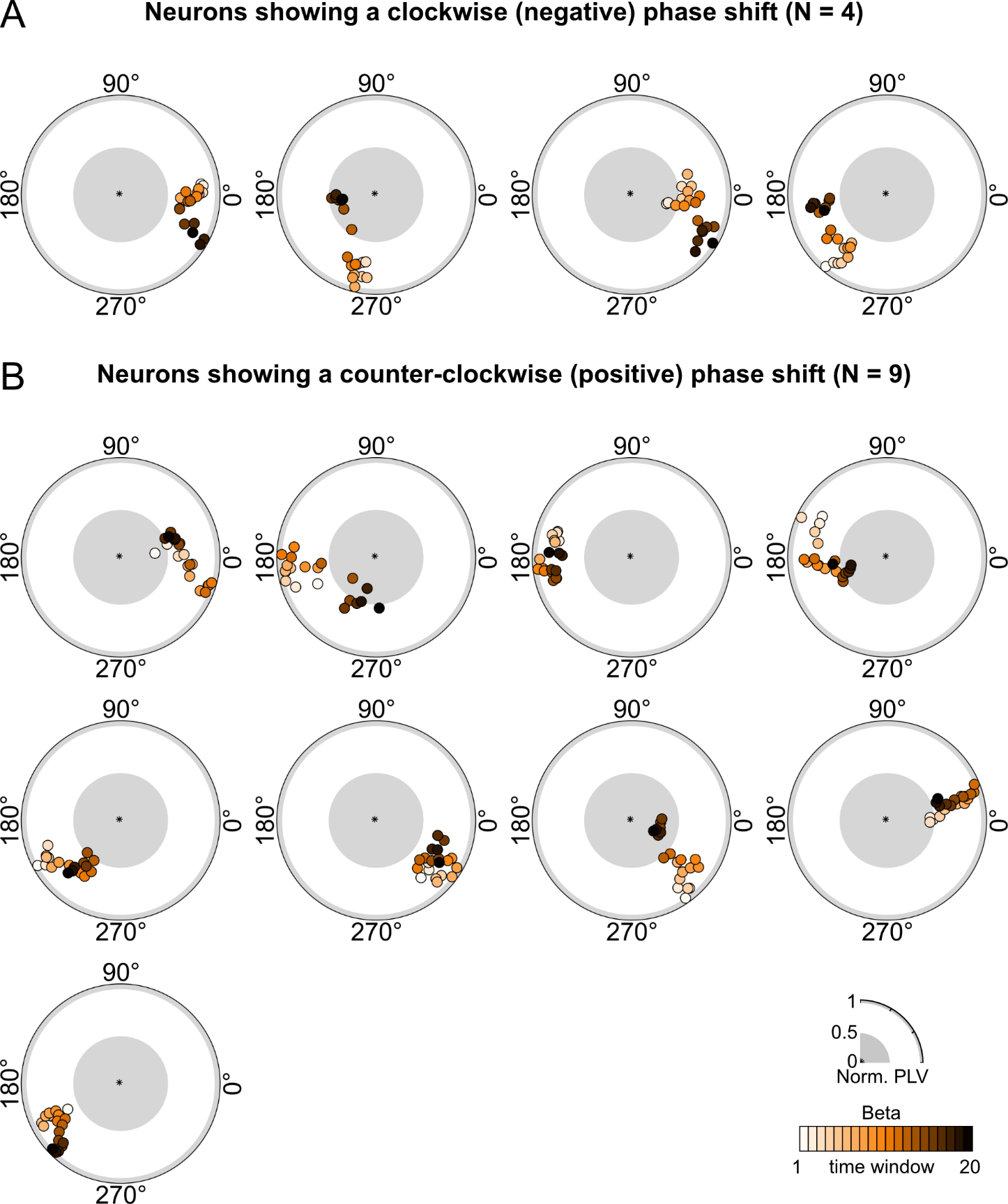

**Supplementary Fig. S18.** Neurons that had a significant circular-linear correlation between time window and preferred phase during beta AC stimulation. Only neurons with a PLV significantly larger than 0 in at least 10 time windows were selected. A) All neurons (N = 4) that showed phase precession in a clockwise direction. B) All neurons (N = 9) that showed phase precession in a counter-clockwise direction.

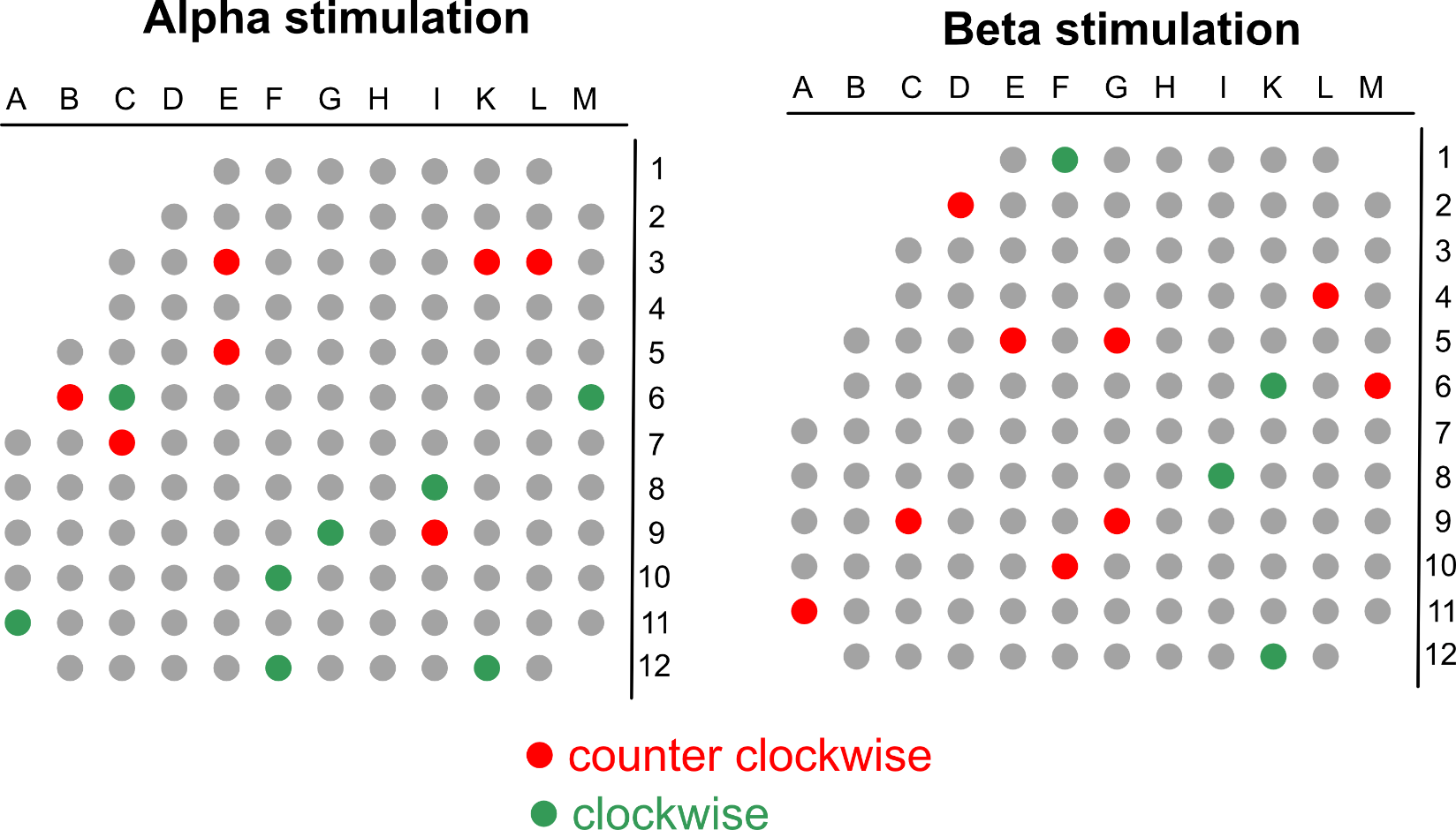

**Supplementary Fig. S19.** Spatial representation of units with neurons showing clockwise (green) and counter-clockwise phase precession, for alpha (left) and beta (right) stimulation.

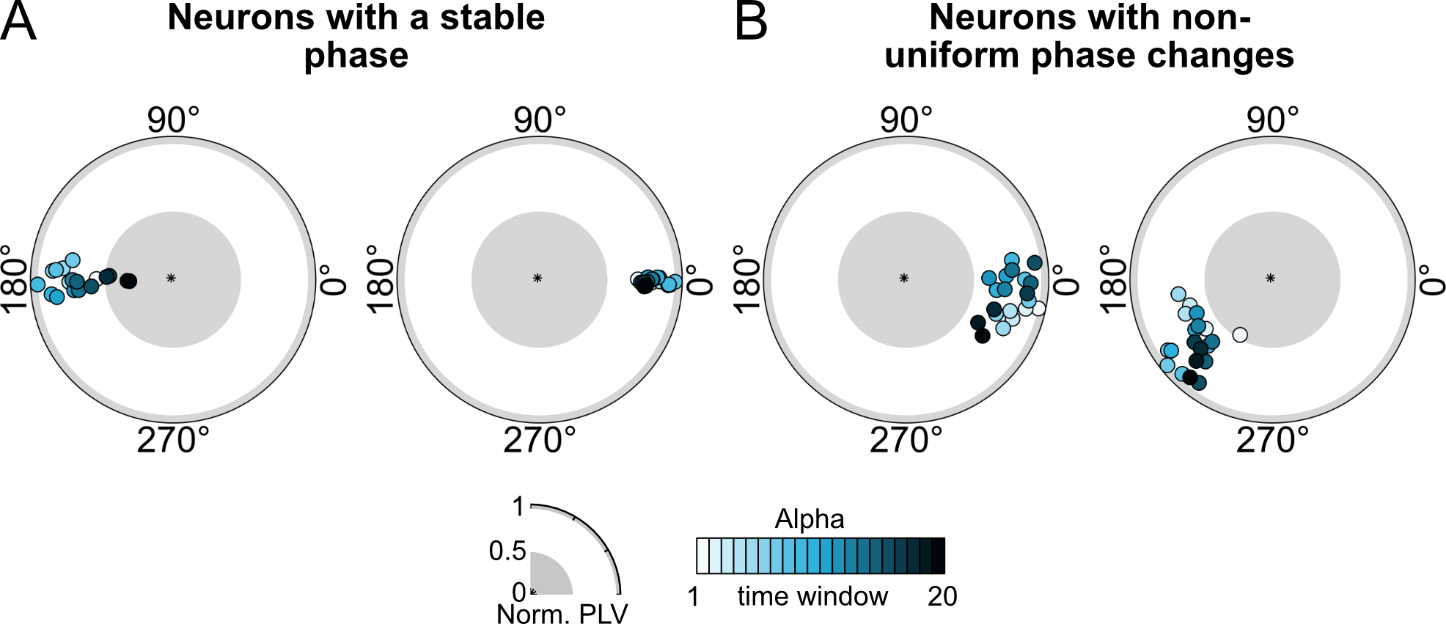

**Supplementary Fig. S20.** A) Two example neurons that showed significant entrainment at a stable phase during alpha AC stimulation. These were defined as neurons that were responsive in ≥10 windows but did have no phase shift ≥15°. B) Two example neurons that showed significant entrainment with non-linear phase shifts during alpha AC stimulation. These were defined as neurons that were responsive in ≥10 windows but did have no significant circular-linear correlation.

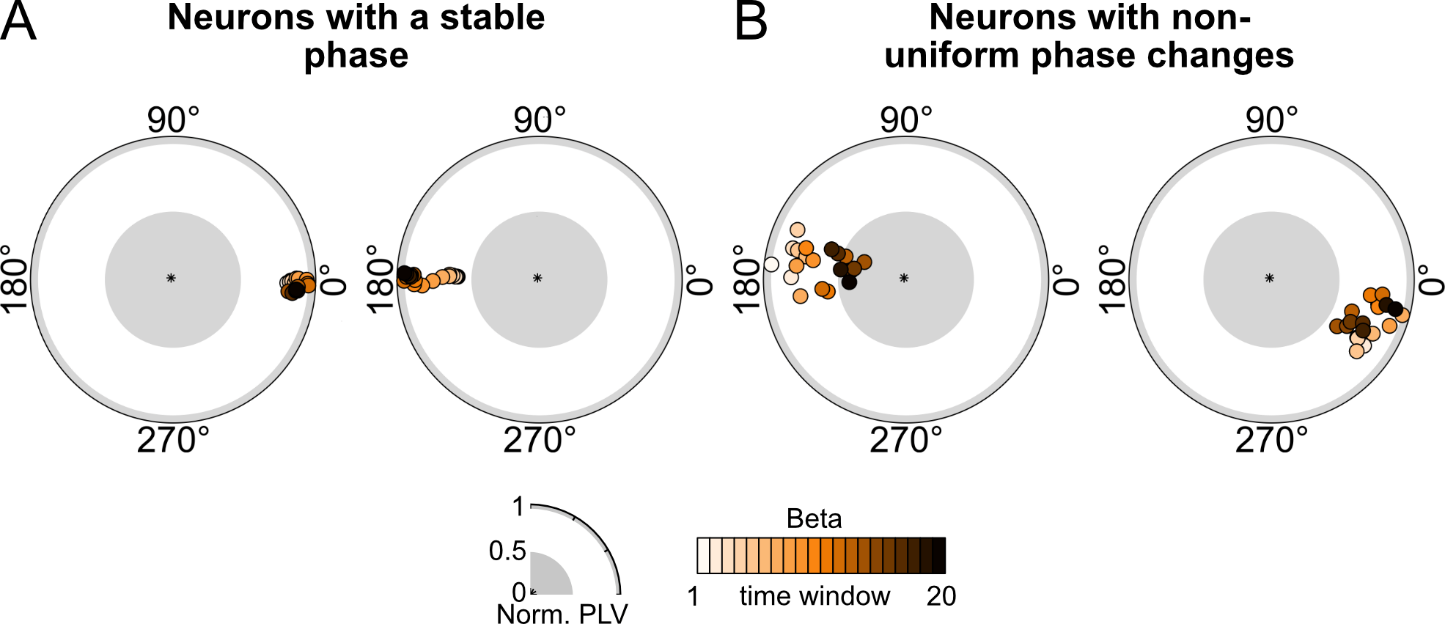

**Supplementary Fig. S21.** A) Two example neurons that showed significant entrainment at a stable phase during beta AC stimulation. These were defined as neurons that were responsive in ≥10 windows but did have no phase shift ≥15°. B) Two example neurons that showed significant entrainment with non-linear phase shifts during beta AC stimulation. These were defined as neurons that were responsive in ≥10 windows but did have no significant circular-linear correlation.

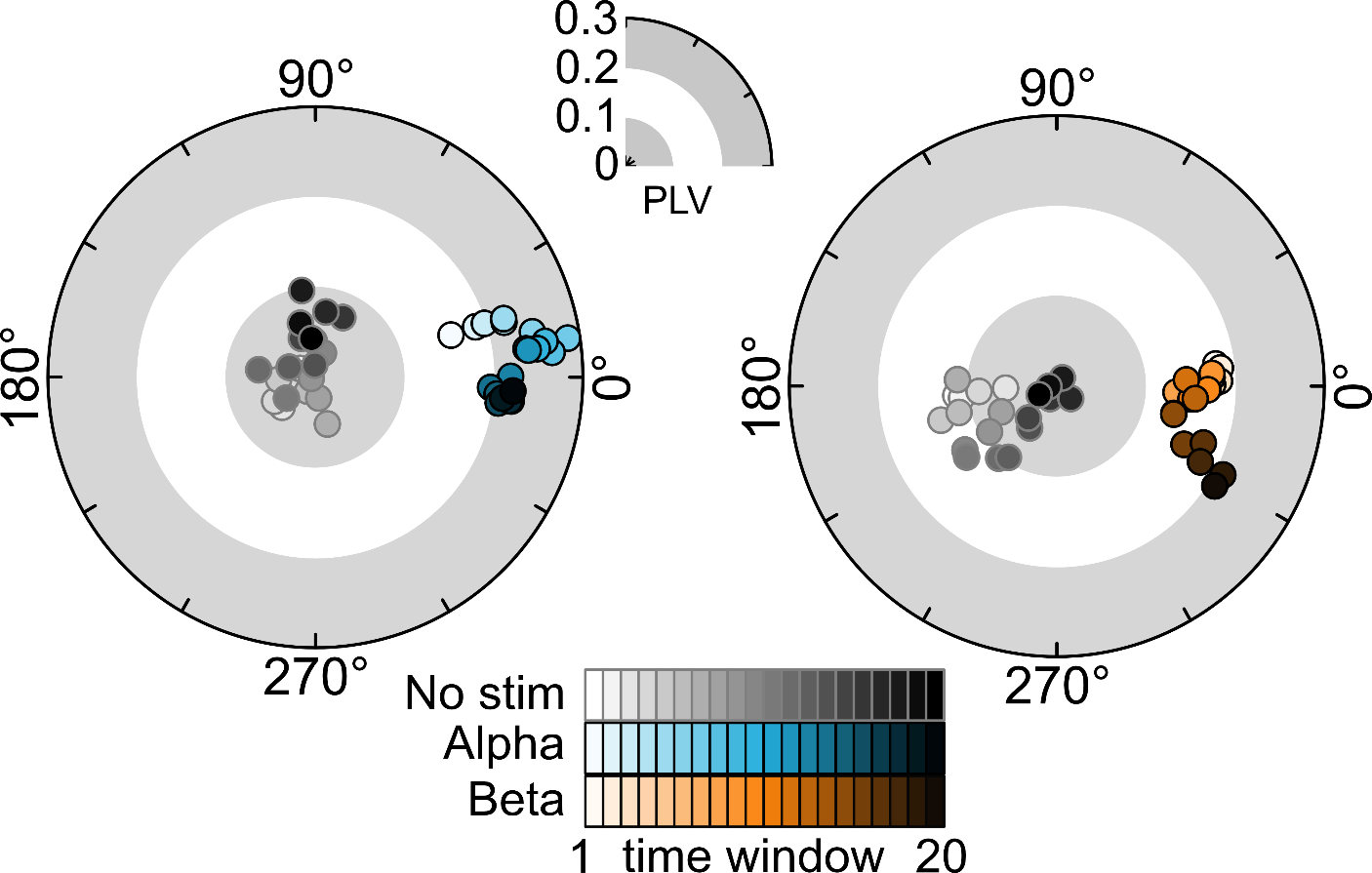

**Supplementary Fig. S22.** Phase shifts during alpha (blue) and beta (orange) stimulation in two example neurons, compared to the same neuron in a 6-minute no stimulation period. During the no stimulation period phase of neural spiking was determined with respect to a virtual tACS signal. The same thresholding steps to determine significance of phase precession (Supplementary Fig. S14). Like the two examples shown above, none of the 28 neurons (alpha N = 15, beta N = 13) that displayed significant phase precession during stimulation showed significant phase shifts during the no stimulation period.

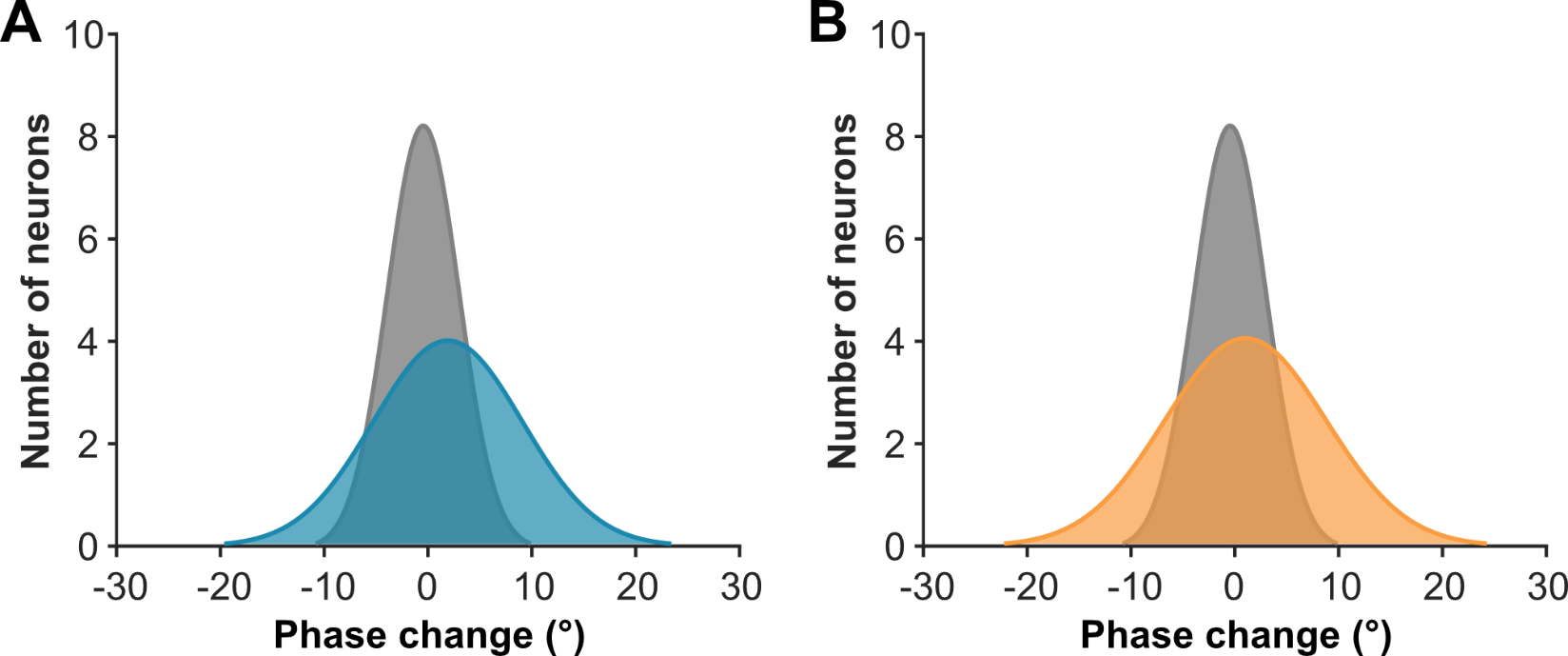

**Supplementary Fig. S23**. Distribution of the average window-to-window phase shift difference for all entrained neurons for alpha tACS (blue, A), beta tACS (orange, B), and no stimulation (gray, A and B). Distributions of phase shifts when no stimulation is applied are quite narrow suggesting quite limited spontaneous phase changes. Alternatively, during tACS distributions are broader, suggesting a wider range of phase shifts induced by tACS.

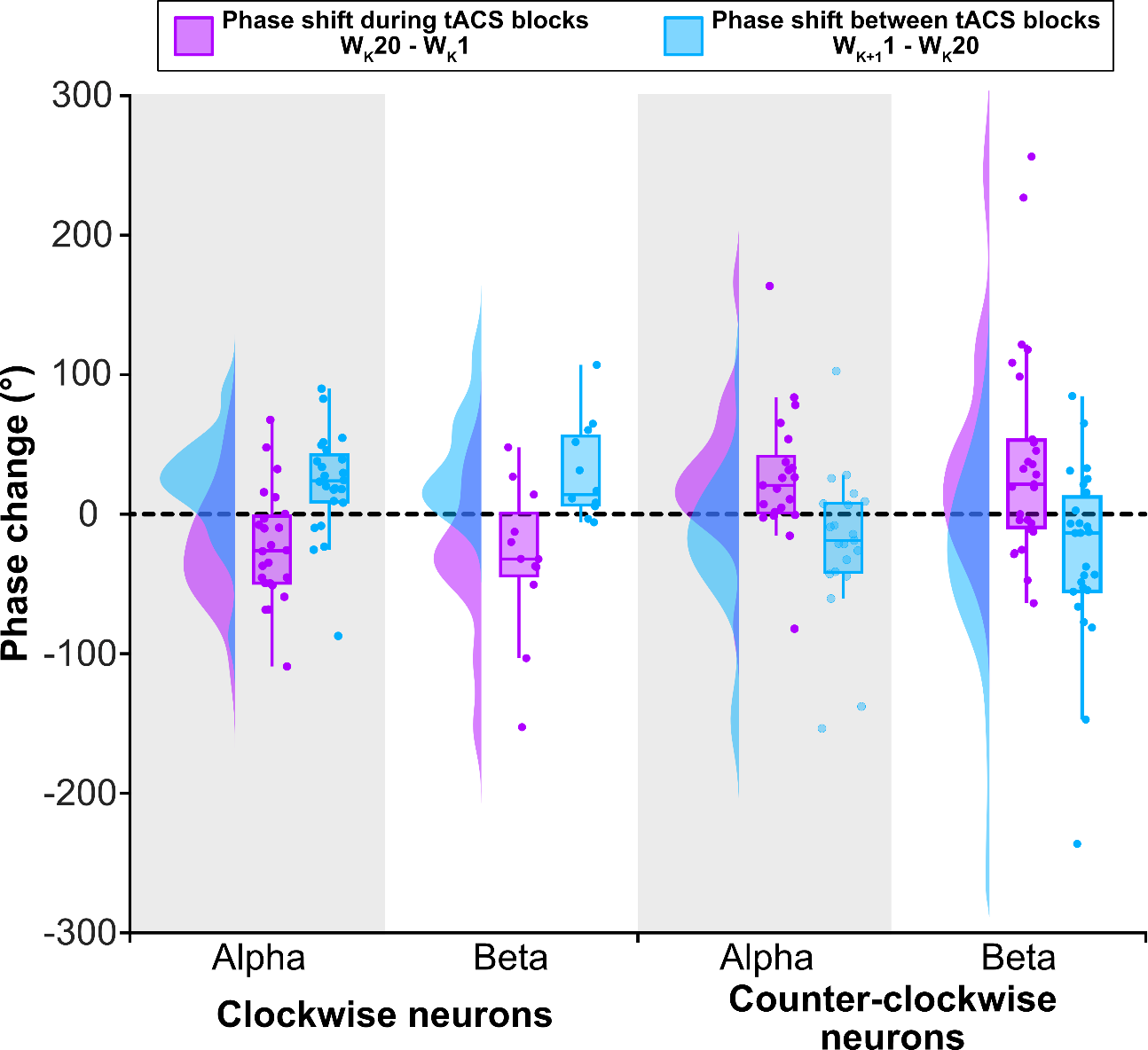

**Supplementary Figure 24**. In purple: Phase change during alpha and beta tACS per block for all clockwise and counter-clockwise neurons. For this the preferred phase at the first (1) window (W) of block k was subtracted from the preferred phase of the last (20) window (W) of block k. Naturally, for neurons that show an overall clockwise phase shift (main text, Figure 3), per block neurons also mostly display a clockwise shift (phase change < 0). Similarly, for neurons that shows an overall counter-clockwise phase shift (main text, Figure 3), per block neurons also mostly display a counter-clockwise shift (phase change > 0). In blue: Phase change between stimulation blocks (at rest). For this the preferred phase of the final (20) window (W) of a block (k) was subtracted from the first (1) window (W) of the subsequent block (k+1). On average phase change between blocks is of similar magnitude but in opposed direction compared to the change during blocks (purple). This suggests a phase reset between blocks, as was also observed in humans (Supplementary Fig. 7).

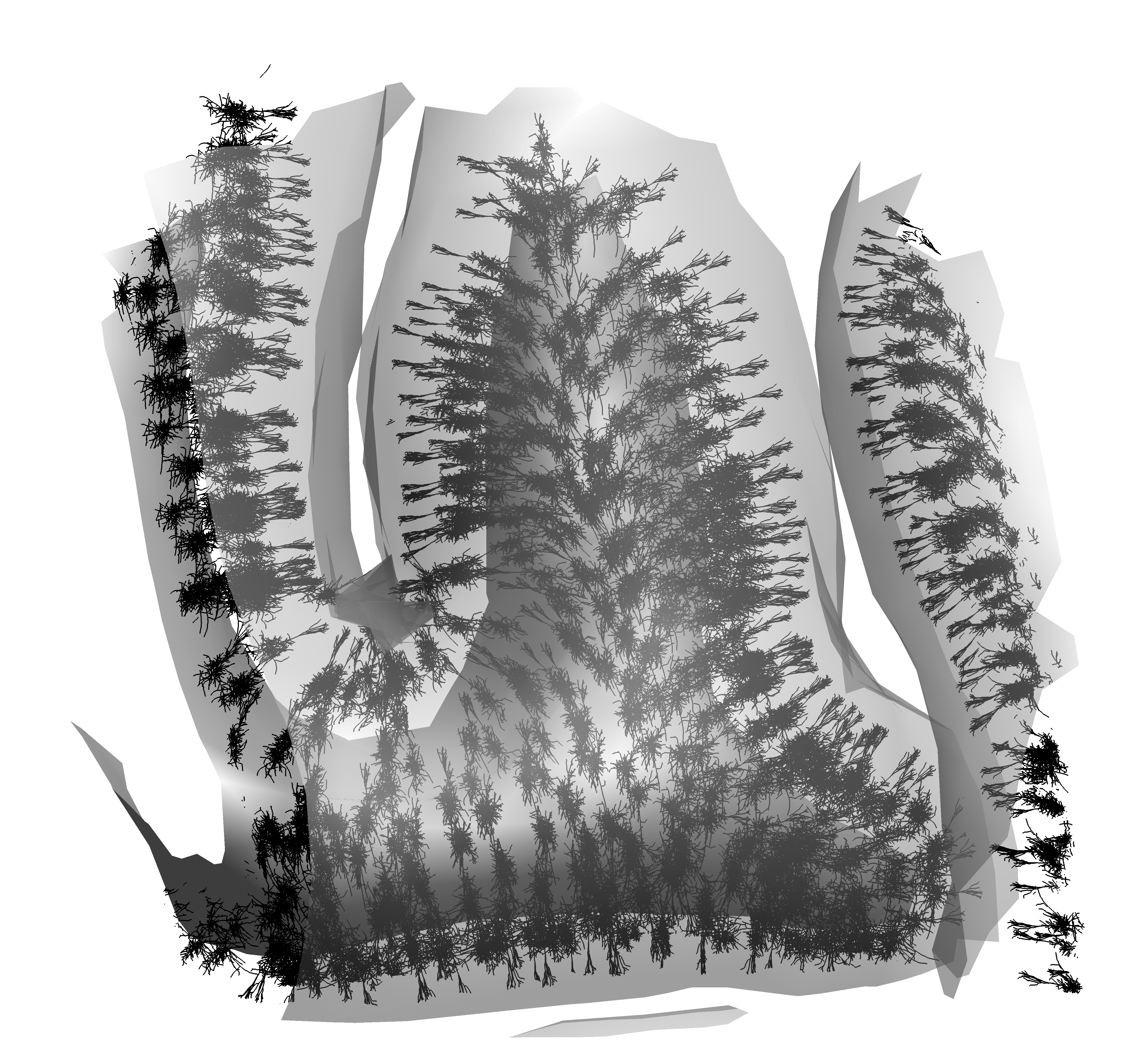

**Supplementary Fig. S25.** The ROI of the precentral gyrus area was populated with 4650 realistic neurons. The gyrus was populated with L5 pyramidal neuron models in 930 tetrahedral elements (5 neuron variants per tetrahedra). The gray matter surface mesh of the ROI contains 930 triangular elements. The soma was placed in the center of each element at a normalized depth of 0.65 between the gray and white matter surface meshes^3^. These model neurons were oriented so that their somatodendritic axes are normal to the gray matter surface.

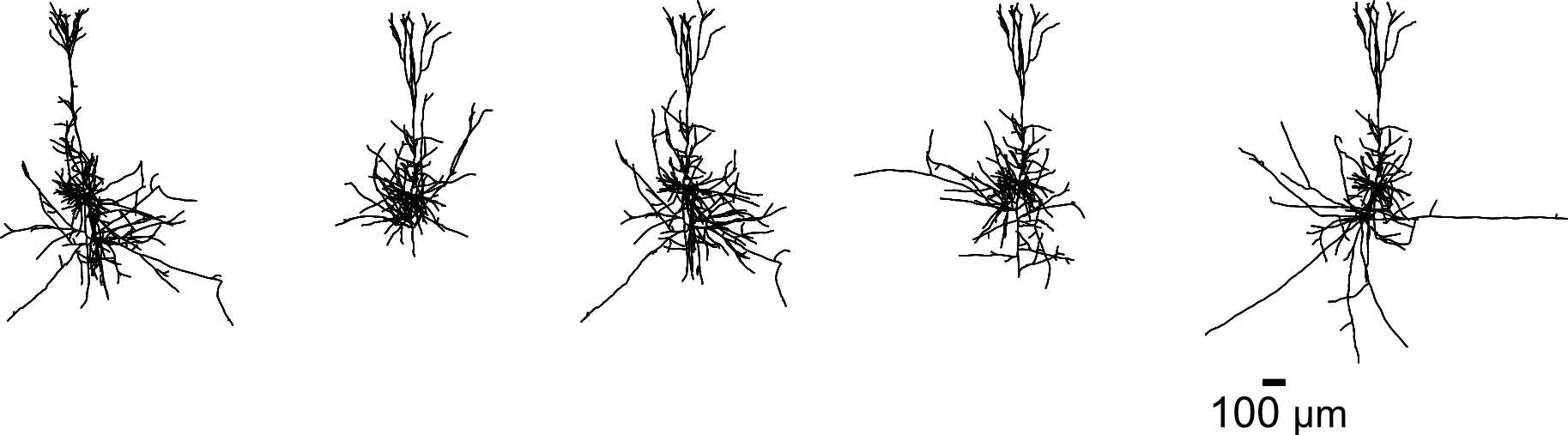

**Supplementary Fig. S26.** The five neuron variants that were used for population of the precentral gyrus ROI^4^.

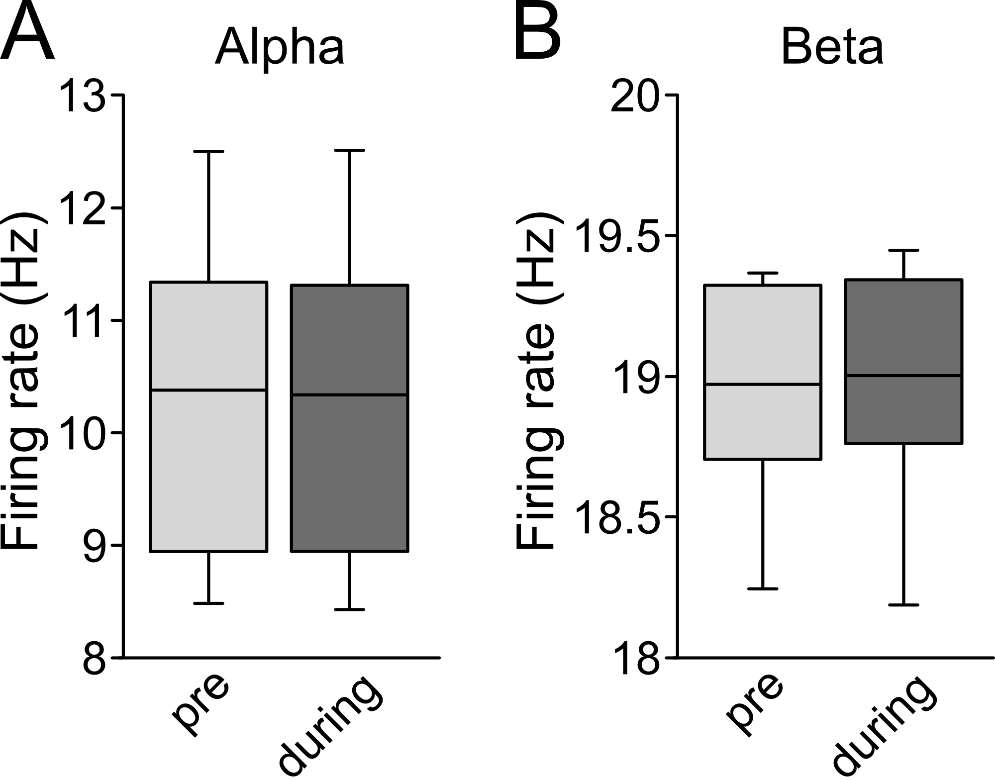

**Supplementary Fig. S27.** Firing rates in the motor cortex strip in a computation model, populated with 4650 realistic neurons. The firing rates before and during AC stimulation were equivalent for both A) alpha stimulation and B) beta stimulation. This suggests that tACS has no effect on firing rate. Note that synaptic inputs were tuned for cells to have similar intrinsic firing rates (mean ± standard deviation, alpha: 10.33 ± 1.49 Hz, beta: 18.92 ± 0.42 Hz).

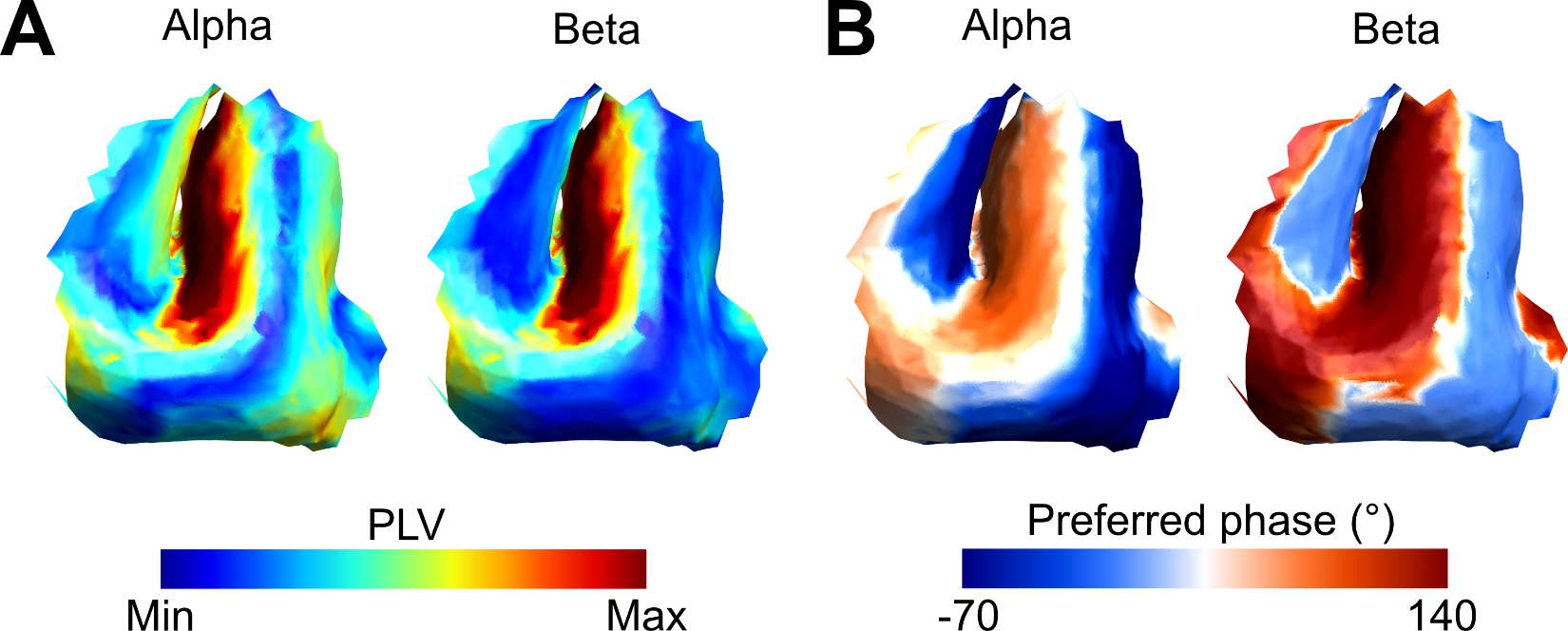

**Supplementary Fig. 28**. Simulations equivalent to those in Fig. 4, but with intrinsic firing rates based on baseline recordings of experiment 2 (3.02 spikes per second and 2.74 spikes per second for the alpha and beta blocks respectively). The results are similar to the original analysis with (A) stronger entrainment in the anterior and posterior wall of the precentral gyrus, compared to the crown. B) Entrainment of the anterior wall (BA6) is more likely at phases between 90 and 180°, whereas entrainment in the posterior wall (BA4) occurs at phases between -90° and 0°.

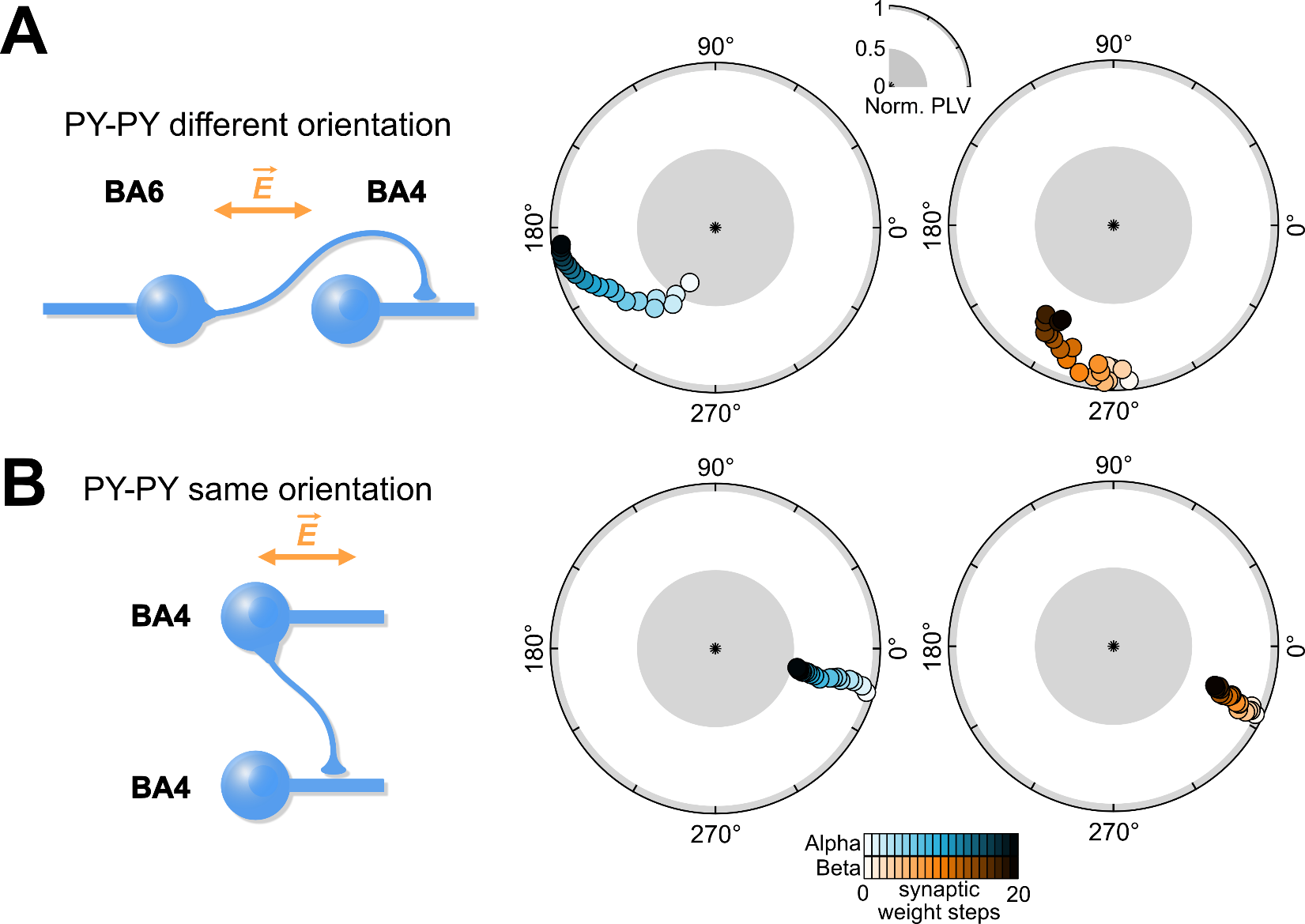

**Supplementary Fig. S29.** We explored the effects of two alternative simplified network models. A) a model with a pyramidal-pyramidal connection with opposing neuron orientations and without inhibitory interneuron. This would reflect an uninhibited connection from BA6 to BA4. In contrast to the original model in the main results, this model showed a clockwise phase shift with increasing synaptic weights, going from ~240° to ~180° for alpha stimulation, and going from ~280° to ~230° for beta stimulation. B) A model with a pyramidal-pyramidal connection with the same neuron orientations and without inhibitory interneuron. This would reflect an uninhibited connection within BA4. The results of this model showed no prominent phase shift for either stimulation frequencies (Fig. 4E).

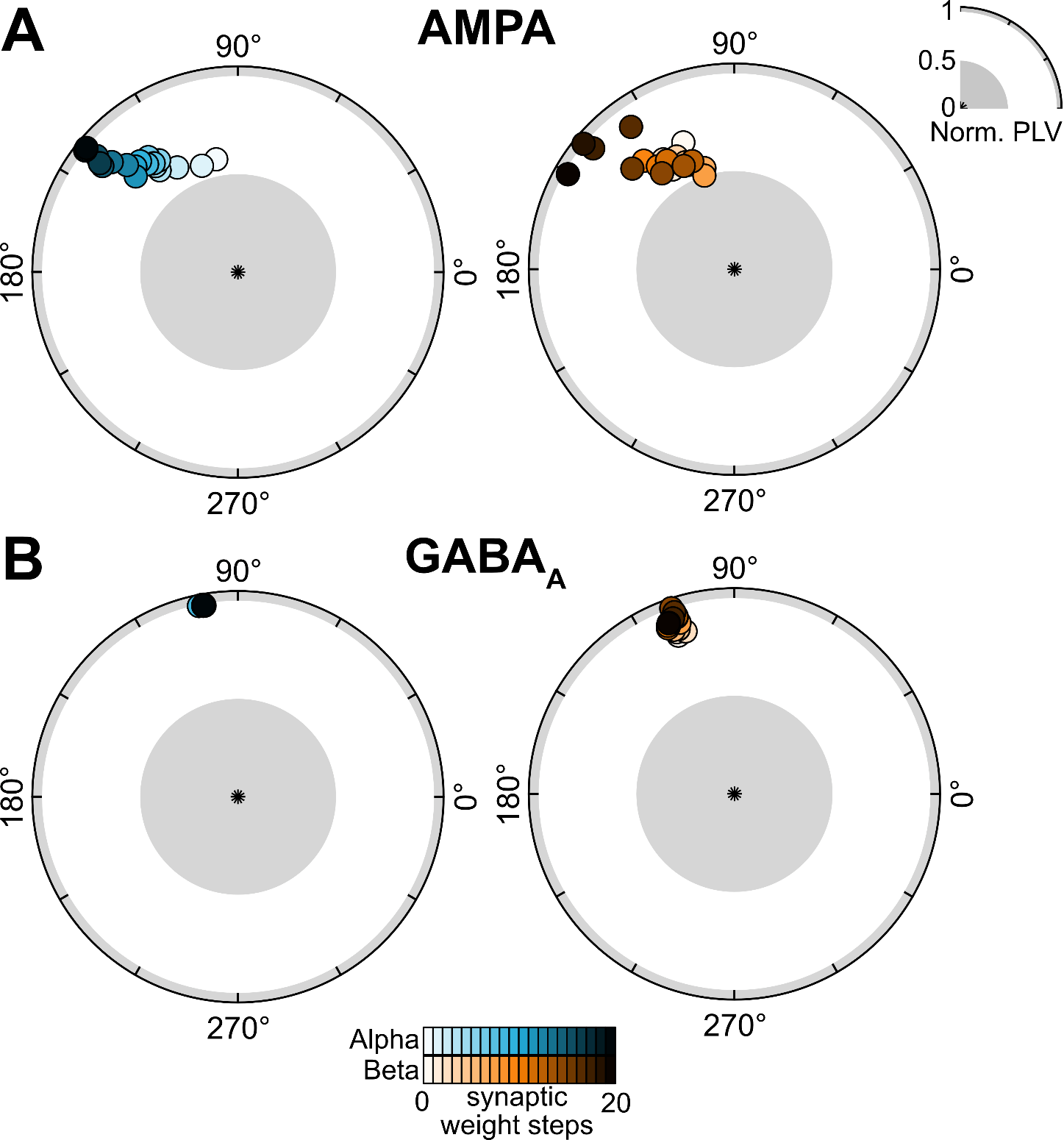

**Supplementary Fig. S30.** Within the original network model (One PY in BA6, one PY in BA4, and one IN in BA4, Fig. 4D) we investigated the contribution of synaptic weight changes of AMPA and GABA_A_ connections to the BA4 PY neuron. A) For AMPA, we used the same synaptic weight step as NMDA. We observed phase shifts that were similar to NMDA, particularly for alpha stimulation (shift from ~90° to ~150°). For beta stimulation also a counter-clockwise phase shift was observed (from ~110° to ~150°), but the trajectory fitted the experimental data less well compared synaptic changes after NMDA. B) For GABA_A_ we did not observe any phase shifts, even if synaptic weights steps were multiplied by 10 compared to AMPA and NMDA. To ensure that GABA_A_ is not associated with phase shifts we multiplied synaptic weights steps for GABA_A_ by 10. This suggests that our experimental data cannot be explained by synaptic changes in GABA_A_.

**Supplementary Table 1.** Overview of alternating current stimulation parameters and experimental design for the three experiments of this study.

|  | **Experiment 1 (human)** | **Experiment 2 (NHP)** | **Experiment 3a (population modelling)** | **Experiment 3b (microcircuit modelling)** |
| --- | --- | --- | --- | --- |
| **tACS intensity** | 1 mA | 1 mA | n/A | n/A |
| **Max. e-field** | 0.31 mV/mm | 0.84 mV/mm | 1.5 mV/mm | 3 mV/mm |
| **tACS duration** | 4 x 6 min (24 min total) | 4 x 6 min (24 min total) | 6 min | 5 min |
| **tACS frequency** | On average: 9.92 Hz and 20.24 Hz | 10 Hz and 20 Hz | 10 Hz and 20 Hz | 10 Hz and 20 Hz |
| **tACS montage** | 2 PiStim (Ø 3.14 cm^2^) electrodes 7 cm anterior and posterior of M1 | 2 PiStim (Ø 3.14 cm^2^) electrodes Fp1 and PO3 | 2 PiStim (Ø 3.14 cm^2^) electrodes 7 cm anterior and posterior of M1 (modelled) | 2 PiStim (Ø 3.14 cm^2^) electrodes 7 cm anterior and posterior of M1 (modelled) |
| **Outcome measure** | MEP | Single-unit activity | Single-unit activity | Single-unit activity |
| **Main finding** | 70.9° (alpha tACS) and 57.4° (beta tACS) phase shift on averaged normalized MEP over 6 min | Phase shifts in 15/46 (alpha tACS) and 13/48 neurons (beta tACS), ranging between ~\|15°\| and ~\|80°\| over 6 min | AC-induced increase in phase locking in anterior and posterior motor cortex depending on electric field direction | ~60° (alpha tACS) and ~80° (beta tACS) phase shift on M1 spiking due to increased NMDA weights between BA6 and BA4 |

| **Supplementary Table 2.** Influence of the shift value on the number of neurons. The default shift value we used in the main manuscript is \|15°\|. The higher the threshold value is, the more selective we are. Even with very high value, few neurons exhibit a positive/negative phase shift. | | | | | |
| --- | --- | --- | --- | --- | --- |
| **Alpha stimulation** | | | | | |
| Shift (°) | \|10\| | **\|15\|** | \|20\| | \|25\| | \|30\| |
| Positive | 10 | 7 | 3 | 3 | 2 |
| Negative | 10 | 8 | 8 | 7 | 5 |
| Total | 20 | 15 | 11 | 10 | 7 |
| **Beta stimulation** | | | | | |
| Shift (°) | \|10\| | **\|15\|** | \|20\| | \|25\| | \|30\| |
| Positive | 14 | 9 | 4 | 3 | 2 |
| Negative | 7 | 4 | 3 | 3 | 3 |
| Total | 21 | 13 | 7 | 6 | 5 |

| **Supplementary Table 3.** Dimensions of neuron compartments (µm) | | | | |
| --- | --- | --- | --- | --- |
|  | Pyramidal neuron | | Interneuron | |
|  | Length | Diameter | Length | Diameter |
| Soma | 30 | 20 | 25 | 25 |
| Dendrite | 2000 | 3.18 | 200 | 3.18 |

| **Supplementary Table 4.** Time constants and reversal potentials for each synapse | | | |
| --- | --- | --- | --- |
| Receptor | $\tau_{1}(ms)$ | $\tau_{2}$(ms) | $E_{syn}$ |
| AMPA | 0.2 | 1.7 | 0 |
| NMDA | 2 | 26 | 0 |
| GABA_A_ | 0.3 | 2.5 | -70 |

| **Supplementary Table 5.** Microcircuit parameters | | | | |
| --- | --- | --- | --- | --- |
| Source | Target | Synapse | Strength (Alpha/Beta) | Delay (ms) |
| PY1 | IN | AMPA | 0.001 | 2 |
| PY1 | IN | NMDA | 0.002/0.001 | 2 |
| PY1 | PY2 | AMPA | 0.008 | 2 |
| PY1 | PY2 | NMDA | 0.0002 | 2 |
| PY2 | IN | AMPA | 0.003 | 1 |
| PY2 | IN | NMDA | 0.0003 | 1 |
| IN | PY2 | GABA_A_ | 0.5 | 1 |
| Background | PY1 | Poisson input | 0.00108/0.0017 | 0 |
| Background | PY2 | Poisson input | 0.00065/0.002 | 0 |
| Background | IN | Poisson Input | 0.00035 | 0 |
